# RALF peptides modulate immune response in the moss *Physcomitrium patens*

**DOI:** 10.1101/2022.10.21.513210

**Authors:** Anna Mamaeva, Irina Lyapina, Andrey Knyazev, Nina Golub, Timur Mollaev, Elena Chudinova, Sergey Elansky, Vladislav V. Babenko, Vladimir A. Veselovsky, Ksenia M. Klimina, Tatiana Gribova, Daria Kharlampieva, Vassili Lazarev, Igor Fesenko

**Affiliations:** Shemyakin and Ovchinnikov Institute of Bioorganic Chemistry, Russian Academy of Sciences, Moscow, Russia; Peoples Friendship University of Russia (RUDN University), 117198, Moscow, Russia; Federal Research and Clinical Center of Physical-Chemical Medicine of Federal Medical Biological Agency, Moscow, Russia; Moscow Institute of Physics and Technology (National Research University), Dolgoprudny, Moscow region, 141701, Russia; Lomonosov Moscow State University, 119991 Moscow, Russia

**Author notes:** These authors contributed equally to this work and share first authorship (Anna Mamaeva, Irina Lyapina). **Correspondence:** Igor Fesenko.

**Keywords:** RALF, peptide, stress response, immunity

## Abstract

RAPID ALKALINIZATION FACTOR (RALFs) are cysteine-rich peptides that regulate multiple physiological processes in plants. This peptide family has considerably expanded during land plant evolution, but the role of ancient RALFs in modulating stress response is unknown. Here, we used the moss *Physcomitrium patens* as a model to gain insight into the role of RALF peptides in coordination of plant growth and stress response in non-vascular plants. The quantitative proteomic analysis revealed concerted downregulation of M6 metalloproteases and some membrane proteins, including those involved in stress response, in *PpRALF1, 2* and *3* knockout (KO) lines. We found that knockout of *PpRALF2* and *PpRALF3* genes resulted in increased resistance to bacterial and fungal phytopathogens - *Pectobacterium carotovorum* and *Fusarium solani*, suggesting the role of these peptides in negative regulation of immune response in *P. patens*. The comparative transcriptome analysis of *PpRALF3* KO and wild type plants under *Fusarium solani* infection showed the clear difference in regulation of genes belonging to phenylpropanoid pathway and associated with cell wall modification and biogenesis between these genotypes. The follow-up analysis revealed the role of PpRALF3 in growth regulation under abiotic and biotic stress regulation, which suggests the role of RALFs in responses to different adverse conditions. Thus, our study sheds light on the function of the previously uncharacterized PpRALF3 peptide and gives a clue to ancestral functions of RALF peptides in plant stress response.

## 1 Introduction

Plants utilize small secreted peptides as important mediators of many processes, from growth and development to response to stress conditions (Olsson et al., 2019). One of such regulators is the conservative 5 kDa peptide family - RALF (Rapid Alkalinization Factor), which are widely present in terrestrial plants (Campbell and Turner, 2017). The mature RALF peptide contains four cysteine amino acid residues which form two S-S bonds (Frederick et al., 2019). RALF peptides are cleaved from an inactive protein precursor and S1P protease is shown to be involved in the cleavage of a mature RALF peptide from a nonfunctional precursor at the conserved dibasic RR site (Srivastava et al., 2009; Stegmann et al., 2017).

The tandem duplication is considered to play a dominant role in the evolution of RALFs and this peptide family has expanded considerably during land plant evolution (Cao and Shi, 2012; Campbell and Turner, 2017). For example, 37 members of this family were found in *Arabidopsis thaliana*, 25 in *Arabidopsis halleri*, 20 in *Zea mays*, but only 3 in *Physcomitrium patens* (Campbell and Turner, 2017; Ginanjar et al., 2022). RALF peptides are diverged into four clades based on mature peptide region sequence features (Campbell and Turner, 2017). RALF peptides from I, II and III clades contain a specific protease cleavage site and conserved YISY motif, which is important for their recognition through receptors (Campbell and Turner, 2017; Xiao et al., 2019). However, RALF peptides from clade IV lack specific cleavage site, in addition, the conserved motif also changes, thus suggesting diverse functions of the representatives from this clade.

Unlike most other plant peptide hormones, RALF peptides bind to membrane-localized receptor-like kinases with malectin-like extracellular domain instead of leucine-rich repeat domain (Franck et al., 2018). Receptors of RALF peptides belong to the *Catharanthus roseus* receptor-like kinase (CrRLK1L) family and include FERONIA (FER), CrRLK1Ls ANXUR1 (ANX1), ANX2, and BUDDHA’S PAPER SEAL (BUPS) 1 and BUPS2 (Ge et al., 2017, 2019). In addition, LRE LIKE GPI-AP1 (LLG1) was proposed to function as a coreceptor for FER in AtRALF1 perception in Arabidopsis (Li et al., 2015). In bryophytes, such as *Marchantia polymorpha* L., *P. patens*, and *Sphagnum fallax* were identified 1, 5 and 7 CrRLK1Ls, respectively (Solis-Miranda et al., 2020). Also, LEUCINE-RICH REPEAT EXTENSINS (LRX) proteins are shown to bind RALF peptides with high affinity (Mecchia et al., 2017; Zhao et al., 2018; Moussu et al., 2020).

The diversity of receptors and co-receptors that can bind different RALF peptides results in their multiple functions and different signaling pathways activation (Abarca et al., 2021). RALF peptides are involved in the regulation of root growth, pollen tube elongation, formation of nitrogen-fixing nodules and inulin accumulation (Murphy and De Smet, 2014; Tena, 2016; Wieghaus et al., 2019). Also, they are involved in the processes of intercellular communication between sporophyte and gametophyte in land plants (Chevalier et al., 2013; Mecchia et al., 2017; Loubert-Hudon et al., 2020). The function of *RALF* genes that have been identified in bryophyte genomes are still being studied. PpRALF1 and PpRALF2 - out of three PpRALF peptides identified in *P. patens* have been shown to promote protonema tip growth and elongation (Ginanjar et al., 2022). Although, the function of PpRALF3 peptide is still unknown.

Besides regulation of growth and development, RALF peptides are also involved in the regulation of responses to abiotic and biotic stresses (Blackburn et al., 2020). For example, AtRALF8 plays a role in simultaneously controlling responses to drought and nematode attack by allegedly regulating cell wall remodeling (Atkinson et al., 2013). However, as has been shown in Arabidopsis, different RALF peptides may have opposite effects. For example, AtRALF17 increases reactive oxygen species (ROS) production and resistance to *Pseudomonas syringae* pv. tomato, while AtRALF23 has the opposite effect (Stegmann et al., 2017). AtRALF33, a close relative of AtRALF23, was shown to inhibit pathogen elicitor-induced ROS production (Stegmann et al., 2017). In addition, AtRALF23 along with AtRALF22 were also shown to participate in regulation of growth and salt stress tolerance by cell wall remodeling (Zhao et al., 2018). The exogenous treatment with synthetic AtRALF1, AtRALF4, AtRALF19 and AtRALF22 reduces elicitor-induced ROS production (Abarca et al., 2021). The corresponding precursors of these AtRALFs contain an S1P cleavage site, but AtRALF6-13, AtRALF15-17, AtRALF20, AtRALF24, AtRALF29-32, AtRALF35-36, which reported to increase elf18-induced ROS production, are not cleaved by this protease (Abarca et al., 2021). Perhaps, the presence of S1P cleavage sites is linked to the negative regulation of the immunity with some exceptions (Stegmann et al., 2017; Abarca et al., 2021).

The participation of RALF peptides in modulation of the response to biotic stress has been shown not only in Arabidopsis, but also in other plants, including crops (Stegmann et al., 2017; Merino et al., 2019). Generally, RALF peptides are considered to be negative regulators of the immune response, which can be used by pathogens to promote infection. Considering that many pathogens, especially fungi, prefer alkaline conditions, the ability of RALFs to alkalize the environment came in very handy. Many plant pathogens synthesize RALF-like peptides that enhance the development of infection (Tena, 2016; Thynne et al., 2017). In tomato and *Nicotiana benthamiana* a synthetic RALF-like peptide from *Fusarium oxysporum* f. sp. *lycopersici* was able to induce ROS burst, alkalinization and activation of MAPKs as well as inhibit the seedlings growth (Thynne et al., 2017). Moreover, RALF-like peptides from plant root-knot nematodes facilitate the process of infection in Arabidopsis and rice (Zhang et al., 2020a). Although these homologs may have an arguable impact on the potency of pathogen infection (Wood et al., 2020).

The role of ancient RALF peptides in response to stress conditions is unknown. Meanwhile, the analysis of different plant lineages allows us to gain insight into their possible functions in regulation of stress response during terrestrialization. Using quantitative proteomic and transcriptomic analysis along with infection with *Pectobacterium carotovorum* and *Fusarium solani*, we studied the role of three PpRALF peptides from the model bryophyte *Physcomitrium patens* in immune stress response. Our study showed the specific role of PpRALF3 in negative regulation of plant stress response.

## 2 Materials and Methods

### 2.1 Plant growth conditions and treatments

*Physcomitrella patens* subsp. *patens* (*Physcomitrium patens* “Gransden 2004”, Freiburg) protonemata were grown on BCD medium supplemented with 5 mM ammonium tartrate (BCDAT) and/or 0.5% glucose with 1.5% agar (Helicon, Moscow, Russian Federation) in a Sanyo Plant Growth Incubator MLR-352H (Panasonic, Osaka, Japan) with a photon flux of 61 μmol/m^2^•s during a 16-hour photoperiod at 24°C in 9-cm Petri dishes (Nishiyama et al. 2000). For proteomic and qRT-PCR analyses, the protonemata were grown in liquid BCDAT medium and collected on day 7. The gametophores were grown on free-ammonium tartrate BCD medium under the same conditions, and 8-week-old gametophores were used for analysis. For resistance to abiotic stresses analyses, protonema was grown on BCD medium without ammonium tartrate supplemented with 150 mM NaCl or 2 mM PQ. For Pp*RALF* genes expression analysis 7-days-old protonema was treated with 100 mM hydrogen peroxide for 2 hours.

For morphological analysis, protonema tissue 2 mm in diameter was planted on 9 cm Petri dishes on BCD and BCDAT media.

For growth rate measurements, photographs were taken at day 30 of subcultivation. Protonemal tissues and cells were photographed using a Microscope Digital Eyepiece DCM-510 attached to a Stemi 305 stereomicroscope.

### 2.2 Generation of knockout mutant lines

*PpRALF1* (Pp3c3_15280V3), *PpRALF2* (Pp3c6_7200V3) and *PpRALF3* (Pp3c25_4180V3) knockout lines were created using the CRISPR/Cas9 system (Collonnier et al., 2017). The coding sequences were used to search for guide sequences preceded by a *S. pyogenes* Cas9 PAM motif (NGG) using the web tool CRISPOR (http://crispor.tefor.net/). The guide sequence closest to the translation start site (ATG) was selected for cloning (Supplementary Table S1 and Figure S1). These sequences were cloned into plasmid pBB (Fesenko et al., 2019), yielding the final complete sgRNA expression cassette. Protoplasts were obtained from protonemata as described previously (Fesenko et al., 2015) and transformed by PEG transformation protocol (Schaefer and Zrÿd, 1997) using mixture of three plasmids 1) one of the pBB plasmid carrying guide RNA expression cassette; 2) pACT-CAS9 carrying CAS9 gene; 3) pBNRF plasmid carrying resistance gene to G418. The plasmids pACT-CAS9 and pBNRF were kindly provided by Dr. Fabien Nogué. Independent knockout mutant lines have been obtained. Double knockout lines were obtained by simultaneous transformation with two plasmids with the corresponding guide RNAs.

### 2.3 DNA and RNA isolation and quantitative reverse transcription PCR (qRT-PCR)

Genomic DNA from gametophores was isolated using a commercial kit (Biolabmix, Russia), according to the manufacturer’s recommendations. Total RNA from gametophores and protonemata was isolated using TRIzol™ Reagent according to the manufacturer’s recommendations. RNA quality and quantity were evaluated using electrophoresis on agarose gel with SYBR Green (Biolabmix, Russia). Total RNA concentration of samples was precisely measured using a Nanodrop™One (Thermo Fisher Scientific, USA).

cDNA was synthesized using the MMLV RT kit (Evrogen, Russian) according to the manufacturer’s recommendations. OligodT primers were used to prepare cDNA from 2 μg total RNA after DNase treatment.

Real-time PCR was performed using the HS-qPCR SYBR Blue (2x) (Biolabmix, Russia) on a LightCycler®96 (Roche, Germany). qPCR was carried out in three biological and three technical replicates. Primers for *PpRALF* genes could be found in Supplementary Table S1.

For qPCR analysis of infection severity, primers were designed for *F. solani* or selected from previous research (Kabir et al., 2020) for *P. carotovorum* to specifically amplify pathogenic DNA from the infected moss plants (Supplementary Table S1).

### 2.4 Pathogen inoculation and treatments

Phytopathogens *Pectobacterium carotovorum subsp. atrosepticum* and *Fusarium solani* were used to infect moss plants. Sterile water-treated plants were used as a control. For bacterial treatment by spraying 15 μl of pre-prepared overnight culture with final concentration 9.5×10^6^ cfu/ml were used. *F. solani* spores were obtained from a solid medium culture and applied by spraying the suspension with final concentration 8.3×10^5^ spores/ml. Petri dishes with infected plants were cultivated at room temperature under standard daylight conditions (16h day/8h night). Infected plants were collected after 7 days and frozen in liquid nitrogen.

### 2.5 Protein extraction and trypsin digestion

Protein extraction and trypsin digestion we conducted as described previously (Faurobert et al., 2007; Fesenko et al., 2021b). iTRAQ labeling (Applied Biosystems, Foster City, CA, USA) was conducted according to the manufacturer’s manual. Proteins were labeled with the iTRAQ tags as follows: Wild type biological replicates – 113, 114, 115 isobaric tags; *PpRALF1, PpRALF2* and *PpRALF3* KO biological replicates – 116, 117, 118 isobaric tags for each mutant line.

### 2.6 LC-MC/MC analysis and protein identification and quantification

The LC–MS/MS analysis was performed as described earlier (Fesenko et al., 2021a). Tandem mass spectra were searched with PEAKS Studio version 8.0 software (Bioinfor Inc., CA, USA) against a custom database containing 32 926 proteins from annotated genes in the latest version of the moss genome v3.3 (Lang et al., 2018), 85 chloroplast proteins and 42 mitochondrial proteins. The search parameters were the following: a fragmentation mass tolerance of 0.05 Da; parent ion tolerance of 10 ppm; fixed modification – carbamidomethylation; variable modifications - oxidation (M), deamidation (NQ), and acetylation (Protein N-term). The results were filtered by a 1% FDR, but with a significance threshold not less than 20 (equivalent is P-value < 0.01). The results were filtered by a 1% false discovery rate (FDR). PEAKS Q was used for iTRAQ quantification. Normalization was performed by averaging the abundance of all peptides. Median values were used for averaging. Although iTRAQ quantification usually underestimates the amount of real fold changes between two samples (Ow et al., 2009), we used a very strict filter for differentially expressed proteins. The threshold for calling a protein differentially abundant was calculated based on an s0 parameter (s0=0.1) as described previously (Schessner et al., 2022) with a permutation test repeated 100 times and Benjamini and Hochberg FDR correction (FDR_BH < 1%).

### 2.7 RNA-seq analysis

DNase treatment was carried out with TURBO DNA-free kit (Thermo Fisher Scientific, Waltham, MA, USA), in volumes of 50 μl. RNA cleanup was performed with the Agencourt RNA Clean XP kit (Beckman Coulter, Brea, USA). The concentration and quality of the total RNA were checked by the Quant-it RiboGreen RNA assay (Thermo Fisher Scientific) and the RNA 6000 Pico chip (Agilent Technologies, Santa Clara, CA, USA), respectively.

RNA libraries were prepared using NEBNext Poly(A) mRNA Magnetic Isolation Module and the NEBNext Ultra II Directional RNA Library Prep Kit (NEB), according to the manufacturer’s protocol. The library underwent a final cleanup using the Agencourt AMPure XP system (Beckman Coulter) after which the libraries’ size distribution and quality were assessed using a high sensitivity DNA chip (Agilent Technologies). Libraries were subsequently quantified by Quant-iT DNA Assay Kit, High Sensitivity (Thermo Fisher Scientific). Finally, equimolar quantities of all libraries (10 pM) were sequenced by a high throughput run on the Illumina HiSeq 2500 using 2 × 100 bp paired-end reads and a 1% Phix spike-in control.

The adaptors and low-quality sequences were removed from raw reads by Trimmomatic v0.39 (Bolger et al., 2014). Clean reads were aligned to the *P. patens* v3.3 reference genome (download from the website: https://phytozome-next.jgi.doe.gov) using HISAT2 v2.1.0 (Kim et al., 2015) and the alignments were sorted with Samtools (Li et al., 2009). The expression abundances of mapped reads were counted by the FeatureCounts tool (Liao et al., 2014). Differential expression analysis was performed by EdgeR package (Robinson et al., 2010). The genes were defined as differentially expressed genes (DEGs) with adjusted p-value ≤ 0.05 and fold change ≥1.0.

### 2.8 Cell wall staining and fluorescence microscopy

Protoplasts were prepared from protonemata as described previously (Fesenko et al. 2015) and incubated for 48 h at solid BCD agar medium. Regenerated protoplasts were dyed with 10 mkg/ml Calcofluor White (Fluorescent Brightener 28) for 5 min. The fluorescence was analyzed using fluorescent microscope (Axio Imager M2, Zeiss, Germany) at λex = 365 nm, BS FT 395, and λem = 445nm /50 nm (Filter set 49 DAPI, Zeiss, Germany).

To detect intracellular ROS, we used the fluorescent dye 2’,7’-Dichlorofluorescin Diacetate (DCFH-DA, Sigma-Aldrich, USA). Seven-days old protonema filaments were harvested with a spatula from the agar surface and placed into mQ. Protonema was then treated with 0.0025% driselase 1 min and incubated with 10 μM DCFH-DA for 15 min in total. The No. 44 filter (λex BP 475 nm/40 nm; λem BP 530 nm/50 nm) was used for DCFH-DA fluorescence detection on the fluorescent microscope Axio Imager M2 (Zeiss) with an AxioCam 506 mono digital camera. Data on the fluorescence intensity were obtained from the related Zeiss software Zen.

### 2.9 Bioinformatic analysis

Protein–protein interaction networks were constructed using STRING v.10 (www.string-db.org) with the default options (Szklarczyk et al., 2014). The visualization of the protein interaction was performed with Cytoscape software (Shannon et al., 2003). The GO enrichment analysis was conducted by g:Profiler (Raudvere et al., 2019). Multiple alignments were created using the MAFFT algorithm (Katoh et al., 2019) and visualized using Jalview software (Waterhouse et al., 2009). Principal Component Analysis (PCA) was performed using iFeature tool (Chen et al., 2018).

### 2.10 Statistics

Statistical analysis and visualization were made in Python v. 3.7.5 (Van Rossum and Drake, 1995) using modules scipy 1.5.2 (Virtanen et al., 2020), seaborn 0.11.1 (Waskom, 2021), numpy 1.20.1, pandas 1.2.3 (McKinney, 2012). For two- or more-way analysis of variance (ANOVA), Tukey’s honestly significant difference (HSD) tests based on multiple comparisons of means were applied to determine which pairwise comparisons were statistically significant. Differences were considered to be significant at p < 0.05.

## 3 Results

### 3.1 The amino acid composition of PpRALFs

The members of a RALF peptide family possess functional heterogeneity in vascular plants (Blackburn et al., 2020), including the role in modulation of immune response. In Arabidopsis, exogenous application of synthetic AtRALF23, AtRALF33, AtRALF34 are shown to inhibit production of reactive oxygen species (ROS) induced upon treatment with immune elicitors (such as elf18) and inhibit seedling and root growth, whereas AtRALF17, AtRALF24, AtRALF32 and some others were able to induce ROS production (Abarca et al., 2021). The previous phylogenetic analysis divided the RALFs into four clades and peptide sequences from different clades could be distinguished based on analysis of their physico-chemical properties (Campbell and Turner, 2017). To test how overall amino acid composition of RALF peptides is related to their possible functions, we used Principal Component Analysis (PCA) to cluster PpRALFs and AtRALFs together. This analysis revealed a clear pattern, in which AtRALF peptides with different effects on pathogen-induced ROS production were separated from each other (Figure 1A). PpRALFs were clustered with AtRALFs that inhibit elicitor-induced ROS production (Abarca et al., 2021). These results are consistent with data of previous phylogenetic analysis (Ginanjar et al., 2022), where PpRALFs were grouped with AtRALF23, AtRALF33, AtRALF23, AtRALF22. These findings suggest that overall amino acid composition of RALF peptides, to some extent, reflect their functional biases and PpRALFs are related to AtRALFs that negatively regulate immune response.

**Figure 1.**
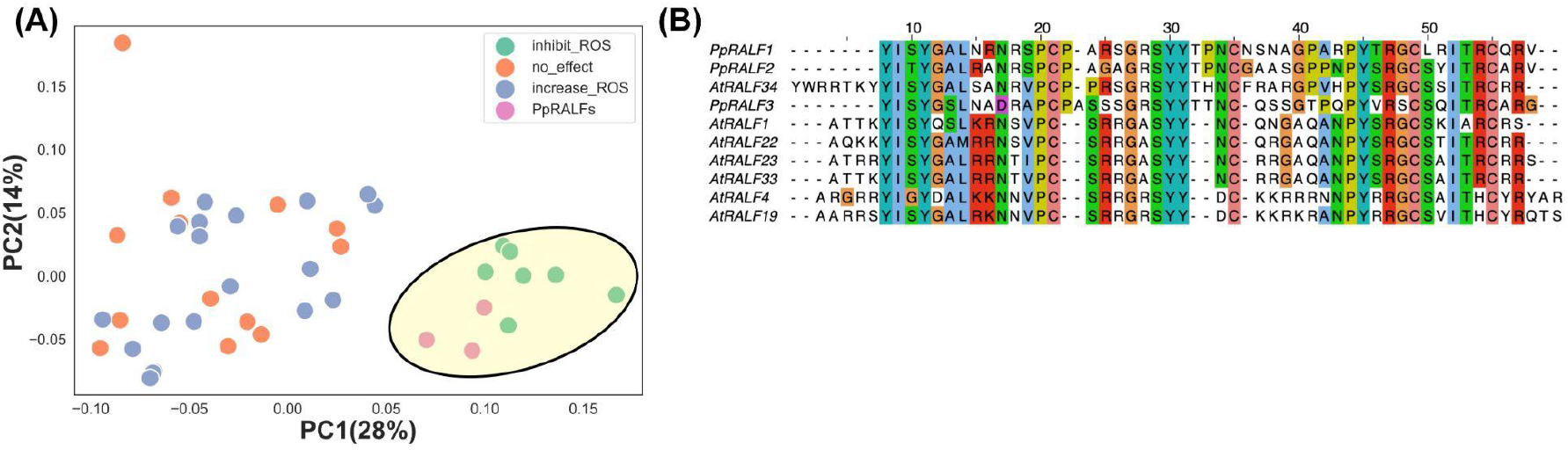
**(A)** - Principal Component Analysis (PCA) of amino acid composition, calculated by iFeature tool (Chen et al., 2018) for AtRALF and PpRALF peptides. Moss PpRALFs were clustered with reported immune-related RALF peptides from Arabidopsis, such as AtRALF1, AtRALF4, AtRALF19, AtRALF22, AtRALF23, AtRALF33, AtRALF34. **(B)** - The multiple pairwise alignment of PpRALFs and immune-related AtRALFs (AtRALF19, AtRALF34, AtRALF33, AtRALF23, AtRALF22).

In *Physcomitrium patens*, PpRALF1 and PpRALF2, that belong to Clade III (Campbell and Turner, 2017), have shown to promote protonema tip growth and elongation (Ginanjar et al., 2022). However, the third member of RALF peptide family - PpRALF3 was not analyzed in Campbell and Turner 2017 and has substitution in conserved “RGC” motif (Figure 1B). The AtRALFs that were clustered together with PpRALF peptides (Figure 1A) belong to Clades 1, 2 and 3 (Campbell and Turner, 2017), implying the absence of the correlation between RALF functions in immune response and division into clades in this case. However, the role of PpRALFs in response to stress conditions is unknown.

### 3.2 The knockout of *PpRALF*s resulted in changes of cell wall-associated proteins

PpRALF1 and PpRALF2 peptides are previously shown to promote tip growth and elongation of protonemata filaments in *P. patens*, but the functions of PpRALF3 peptide are currently unknown (Ginanjar et al., 2022). The chemical synthesis of RALF peptides, as well as the generation of recombinant peptides, is associated with certain difficulties. Proper bonding and folding of peptides require the right conditions, otherwise peptides may not work the way they do, or a higher concentration could be required (Abarca et al., 2021). Since this may not reflect the actual effect of the peptides, we decided to use only knockout lines. At least two independent knockout lines for each *PpRALF* gene and double knockout lines - *PpRALF1* and *PpRALF3*; *PpRALF2* and *PpRALF3* were generated by CRISPR/Cas9 technology (Table 1). However, we failed to obtain triple and double *PpRALF1* and *PpRALF2* knockout lines after several attempts. Probably, this is due to their important roles in regulation of moss growth and development.

**Table 1.** The list of obtained mutant lines.

| Phytozome ID | Name | The number of mutant lines |
| --- | --- | --- |
| Pp3c3_15280 | *PpRALF1 KO* | 2 |
| Pp3c6_7200 | *PpRALF2 KO* | 2 |
| Pp3c25_4180 | *PpRALF3 KO* | 2 |
| Pp3c3_15280, Pp3c25_4180 | *PpRALF1,3 KO* | 1 |
| Pp3c6_7200, Pp3c25_4180 | *PpRALF2,3 KO* | 1 |

It has been previously shown that knockout of *PpRALF* genes results in reducing the formation of caulonema filaments (Ginanjar et al., 2022). Therefore, we tested the growth of our mutants on medium with and without ammonium tartrate (BCDAT/BCD; Figure 2A). The ammonium tartrate reduces the transition into caulonemal cells and promotes chloronemal branching (Vidali and Bezanilla, 2012). The morphology of wild type and single knockout mutants was similar, but the diameter of *PpRALF3* KO plants was significantly larger than wild types on the BCDAT medium (Figure 2A and 2B; ANOVA with post-hoc Tukey HSD *P* < 0.001). In contrast, the growth rate of *PpRALF1* KO lines did not differ from wild type on all media examined (Figure 2A and 2B). In addition, *PpRALF2* KO plants were significantly smaller than wild type plants on the medium with ammonium tartrate (Figure 2B, ANOVA with post-hoc Tukey HSD *P* < 0.001). Both double knockouts showed significant inhibition of the growth rate on medium with and without ammonium tartrate (Figure 2A and 2B; ANOVA with post-hoc Tukey HSD *P* < 0.001). In addition, the gametophores were significantly longer in both *PpRALF3* KO lines in comparison to other genotypes (Supplementary Figure S2; ANOVA with post-hoc Tukey HSD *P* < 0.001). Thus, our results are in line with previously obtained data on morphology and phenotypes of *PpRALF* knockouts (Ginanjar et al., 2022). In addition, our results suggest that all PpRALFs are functional and might play different roles, beyond the regulation of only growth processes.

**Figure 2.**
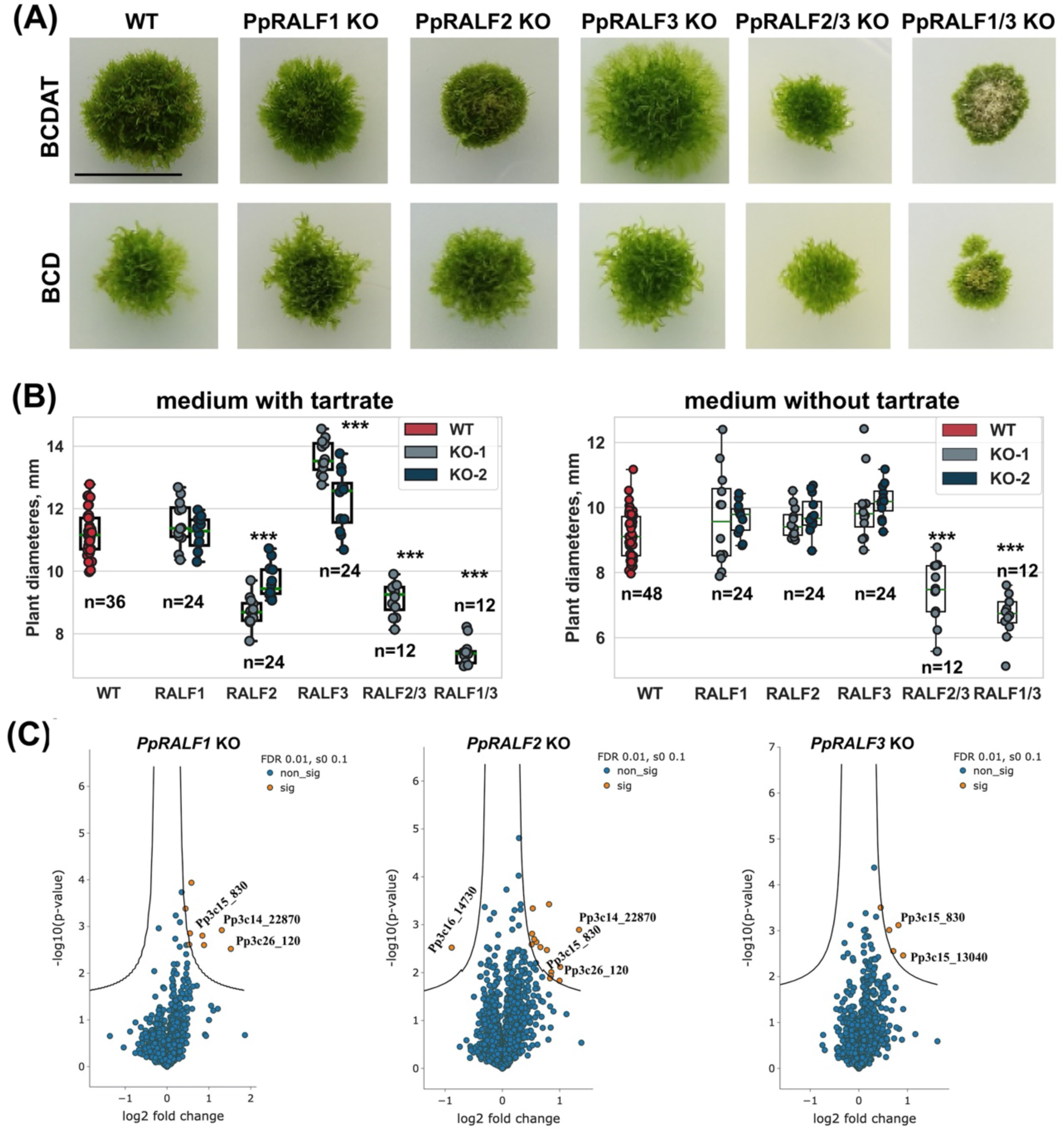
**(A)** - Phenotype of 30-d-old plants of wild type and the obtained knockout lines grown on two mediums with and without ammonium tartrate; **(B)** - The measured diameter of wild type and knockout lines grown on mediums with (left) and without ammonium tartrate (right). Analysis of variance and Tukey’s HSD post hoc tests were performed \*\*\**P* < 0.001; **(C)** - Volcano plots of the entire set of proteins quantified during iTRAQ analysis of *PpRALF1, PpRALF2* and *PpRALF3* KO lines, respectively. Proteins significantly changed in abundance are depicted in orange; Pp3c15_830 - M6 family metalloprotease domain-containing protein, Pp3c14_22870 - Hydrophobic surface binding protein (HsbA), Pp3c26_120 - bryoporin, Pp3c15_13040 - 12-lipoxygenase, Pp3c16_14730 - Stigma-specific protein, Stig1.

Using isobaric tags for relative and absolute quantification (iTRAQ), we further compared proteomes of *PpRALF1, PpRALF2, PpRALF3* knockout lines and wild type plants. In total, we identified 2723 protein groups in *PpRALF1* KO, 3074 protein groups in *PpRALF2* KO and 3078 protein groups in *PpRALF3* KO (Supplementary Table S2). Although iTRAQ quantification usually underestimates the amount of real fold changes between two samples (Ow et al., 2009), we used a very strict cut-off to reliably identify differentially expressed protein groups (DEPs), such as an s0 parameter equal 0.1 and multiple hypothesis correction that was performed based on a permutation test (see Methods). In contrast to previously published transcriptomes of the *PpRALFs* knockout lines (Ginanjar et al., 2022), we have not identified substantial changes in knockout proteomes in comparison to wild type plants, suggesting primarily regulatory roles of PpRALF peptides. In *PpRALF1* KO plants only 8 protein groups were significantly changed (FDR BH < 1%, s0=0.1; Figure 2C; Supplementary Table S2) in comparison to wild type plants and all of these DEPs were downregulated. The knockout of PpRALF2 peptide resulted in downregulation of 15 protein groups (FDR_BH < 1%, s0=0.1; Figure 2C; Supplementary Table S2) and upregulation of an uncharacterized 257 aa protein Pp3c16_14730, containing predicted signal peptide. In *PpRALF3* KO plants only 5 protein groups were significantly downregulated (FDR_BH < 1%, s0=0.1; Figure 2C; Supplementary Table S2).

We further compared these DEPs from all knockout lines and found a core set of four proteins that were changed in at least two single-gene knockout lines. For example, we revealed that a protein Pp3c15_830 (M6 family metalloprotease domain-containing protein) was downregulated in *PpRALF1, PpRALF2* and *PpRALF3* KO lines. However, the role of such proteins in RALF signaling has not been previously described. Another DE protein that belonged to M6 metalloproteases - Pp3c13_22390 was downregulated only in the *PpRALF1* KO line. We also identified two proteins that were downregulated in both *PpRALF1* and *PpRALF2* KO lines. One of them - Pp3c26_120 (Bryoporin) is known to be important for a response to osmotic stress in mosses (Hoang et al., 2009). The other uncharacterized protein - Pp3c14_22870 has orthologs that were characterized as Hydrophobic surface binding proteins (HsbA) in other moss species. We also identified LOX3 protein (Pp3c15_13040) that downregulated in *PpRALF2* and *PpRALF3* KO lines, which might be involved in the arachidonic acid metabolism, suggesting the possible role of these paralogs in biotic stress response.

Thus, the knockout of *PpRALF* genes resulted in downregulation of some proteases and previously uncharacterized membrane proteins. To determine whether these changes in proteomes of mutant lines affected the cell wall regeneration processes, we explored the process of protoplast regeneration in the knockout and wild type genotypes. The protoplasts were dyed with Calcofluor White fluorescent dye that is widely used to visualize cellulose, callose, and other β-glucans in the plant cell wall (Maksimov et al., 2016; Herrera-Ubaldo and de Folter, 2018). On average, the cell wall regenerated 80% faster in *PpRALF2* KO lines (chi-square *P* < 0.001) and 46% faster in *PpRALF3* KO lines (chi-square *P* < 0.001) than in wild type cells after two days of regeneration process (Supplementary Figure S3). We found no significant differences in the cell wall regeneration rate between *PpRALF1* KO-1 and wild type plants.

### 3.3 The *PpRALF3* knockout lines are more tolerant to abiotic stress factors

Some members of the RALF peptide family are shown to modulate the abiotic stress response in angiosperms (Zhao et al., 2018). Therefore, we next sought to expand our understanding of PpRALFs functions under abiotic stress conditions. At first, we used previously obtained RNA-seq data (Khraiwesh et al., 2015) to analyze the expression of *PpRALF* genes under salt, drought, and cold treatments. According to this study, *PpRALF1* was significantly upregulated in all stress conditions after 0.5 h and under cold treatment after 4h but downregulated under drought and salt treatment after 4 h. In addition, the *PpRALF2* gene was significantly upregulated at drought after 0.5 h and at cold after 4 h as well. The *PpRALF3* transcripts were detected only under salt and drought treatment, suggesting its possible role in stress response.

We also assessed the transcriptional level of all *PpRALFs* under hydrogen peroxide treatment for 2 hours. According to qRT-PCR, all *PpRALFs* were significantly downregulated under this treatment (Figure 3A; ANOVA with post-hoc Tukey HSD *P* < 0.001). However, the transcriptional level of *PpRALF3* decreased less than that of other *PpRALFs*.

**Figure 3.**
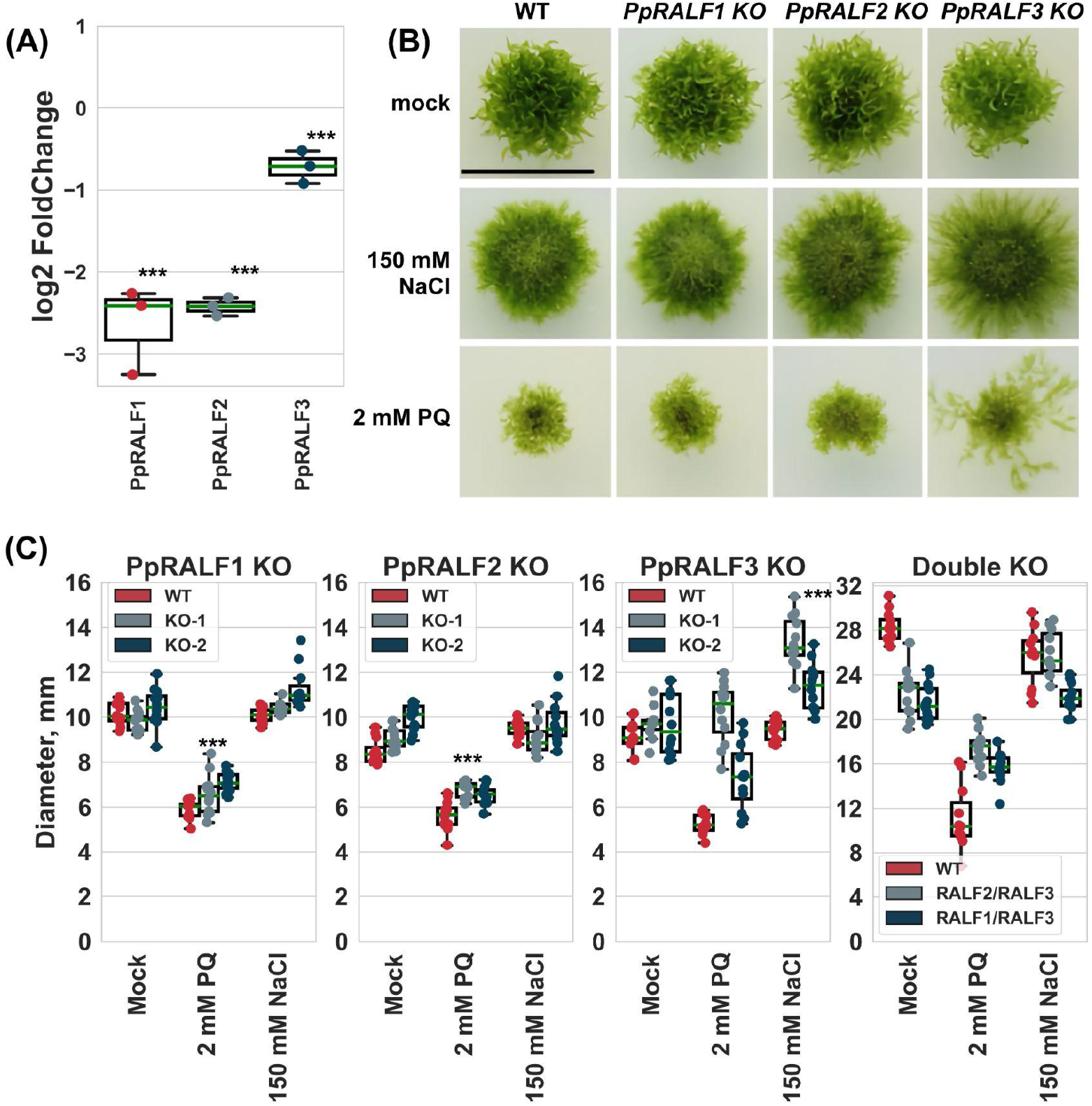
**(A)** - The results of quantitative reverse transcription PCR (RT-qPCR) analysis of the transcriptional level of *PpRALFs* under hydrogen peroxide treatment for 2 hours. Analysis of variance and Tukey’s HSD post hoc tests were performed \*\*\**P* < 0.001; **(B)** - Phenotype of 30-d-old plants of wild type and the knockout lines grown on medium supplemented with 150 mM NaCl or 2 mM paraquat; **(C)** - The measured diameter of moss plants grown on medium supplemented with 150 mM NaCl or 2 mM paraquat. Analysis of variance and Tukey’s HSD post hoc tests were performed \*\*\**P* < 0.001.

Taking into account the role of RALF peptides, such as AtRALF22/23, in regulation of salt stress tolerance in Arabidopsis (Zhao et al., 2018), we further assessed the growth rate of *PpRALF* KO plants upon the oxidative (2 mM paraquat) and salt (150 mM NaCl) treatments. We have chosen the salt concentration of 150 mM NaCl for these experiments based on our assessment of wild type plant phenotypes upon different salt concentrations (Supplementary Figure S4). Our experiments showed that wild type and knockout plants were affected by 150 mM NaCl, confirming the adverse effect of high salt concentration on moss growth and development (Figure 3B). Herewith, the growth rate of wild type and *PpRALF1* and *2* KO plants was similar under salt treatment, except for *PpRALF3* KO lines. The diameter of *PpRALF3* KO lines was significantly larger (ANOVA with post-hoc Tukey HSD *P* < 0.001) under salt treatments compared to mock treated and wild type plants due to diffusion of protonemal filaments (Figure 3B and C). These findings suggest the role of PpRALF3 in the response to abiotic stress factors, as has been shown for angiosperm RALFs. For example, overexpression of AtRALF22 or AtRALF23 has previously been shown to increase sensitivity to salt stress (Zhao et al., 2018).

Under the oxidative stress conditions, the growth and morphology of both wild type and mutant plants were severely affected (Figure 3B). The growth rate of *PpRALF1* and *PpRALF2* KO lines was also significantly suppressed in comparison to normal conditions (Figure 3B and C; ANOVA with post-hoc Tukey HSD *P* < 0.001). However, only the growth of *PpRALF3* KO-2 plants was significantly reduced in comparison to normal conditions (Figure 3C; ANOVA with post-hoc Tukey HSD *P* < 0.001) and both *PpRALF3* mutant lines grew better than wild type. In addition, both double KO mutant lines were less sensitive to paraquat in comparison to wild type plants (Figure 3C). Thus, our results showed that *PpRALF3* is responsive to adverse conditions and its knockout lines are more tolerant to growth inhibition during abiotic stress response.

### 3.4 The knockout of *PpRALF2* and *PpRALF3* increases resistance to phytopathogens

It has been previously shown that some RALF peptides can modulate immune response in vascular plants (Stegmann et al., 2017; Zhang et al., 2020b; Abarca et al., 2021). For example, treatment with AtRALF peptides increased or reduced ROS production under induction of immune response by elf18 (Abarca et al., 2021). To determine whether the knockout of *PpRALF* genes interfere with immune response in bryophytes, we next analyzed ROS production in *P. patens* wild type and knockout lines under driselase treatment. We found that ROS production was significantly reduced in *PpRALF1* and *PpRALF2* KOs in comparison to wild type plants and *PpRALF3* knockout line after driselase treatment (Figure 4A; ANOVA with post-hoc Tukey HSD *P* < 0.001). Moreover, the background level of ROS production in the *PpRALF3* knockout lines without treatment was significantly higher than in the other genotypes (Figure 4A; ANOVA with post-hoc Tukey HSD *P* < 0.001). That may be explained by the fact that PpRALF3 plays a role in suppressing the immune response. This is in concordance with our PCA analysis where PpRALFs tend to cluster with AtRALFs that decrease pathogen-induced ROS production (Figure 1A).

**Figure 4.**
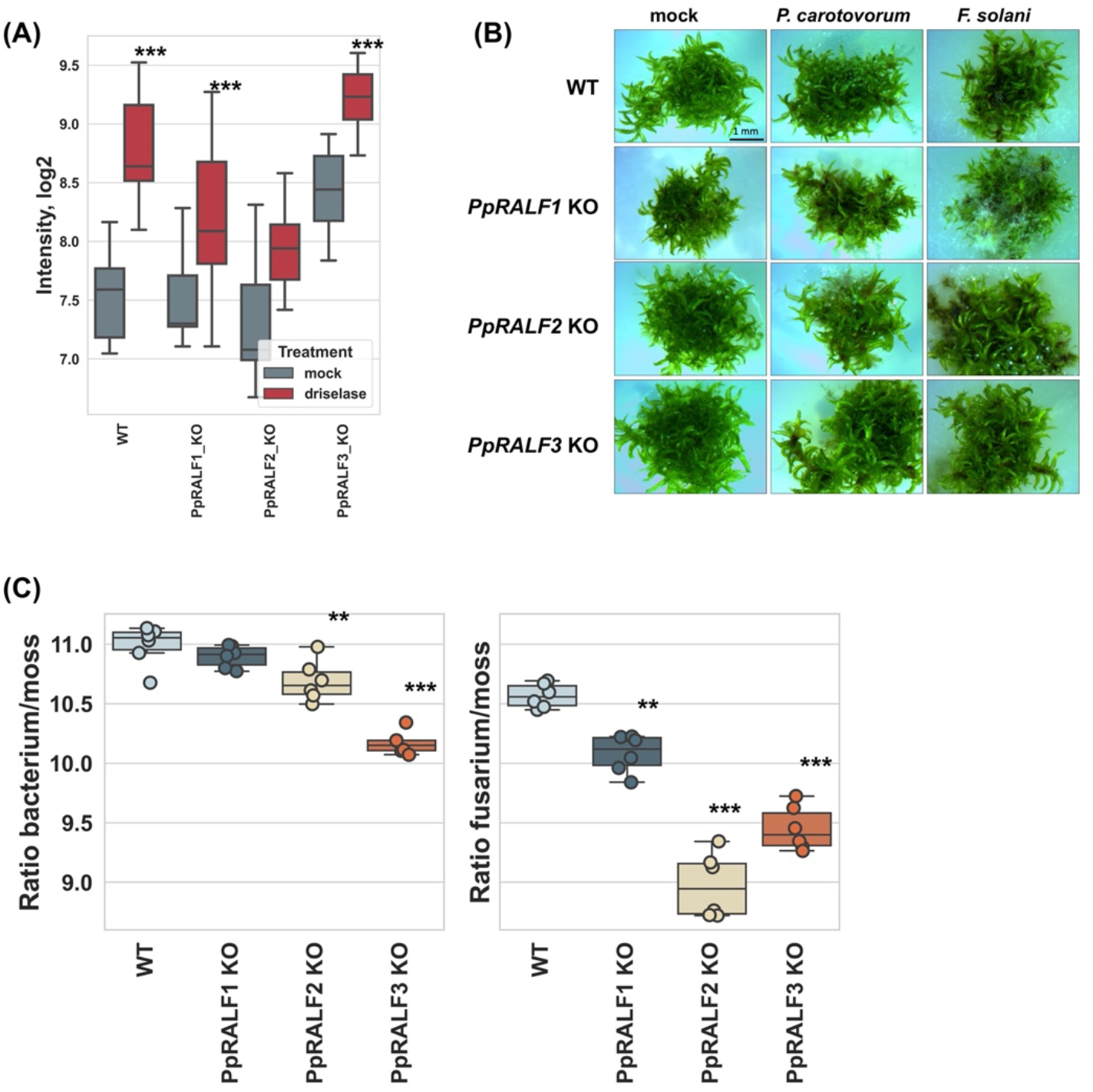
(**A)** - Effect of PpRALF peptide knockouts on ROS production in wild type and knockout lines under treatment with immune elicitors; **(B)** - Symptom development at 7 dpi in wild type, *PpRALF1, PpRALF2* and *PpRALF3* KO moss plants; **(C)** - Ratios of *P. carotovorum* and *F. solani* DNA levels to *P. patens* genomic DNA that were estimated by qPCR analysis. The results of six independent experiments are shown. Analysis of variance and Tukey’s HSD post hoc tests were performed for qPCR data. \*\**P* < 0.01, \*\*\**P* < 0.001.

We further studied the possible functions of PpRALFs in defense response against phytopathogens. For this, we infected moss plants with two well-known phytopathogens - *Pectobacterium carotovorum subsp. atrosepticum* and *Fusarium solani*, which are mildly aggressive and keep plants alive (Figure 4B). These pathogens also show distinct symptoms of infection, such as brown spots caused by tissue necrosis (Akita et al., 2011; Alvarez et al., 2016). To test whether the colonization and growth of *P. carotovorum* and *F. solani* differ between wild type and knockout lines, moss plants were collected at 7 days post inoculation (dpi) and the ratio of pathogen DNA to plant DNA concentrations was measured using quantitative PCR analysis as was described previously (Castro et al., 2016).

According to our experiments, *P. carotovorum* growth rate was significantly higher in wild type plants compared to *PpRALF2* KO and *PpRALF3* KO mutant lines (Figure 4C; ANOVA with post-hoc Tukey HSD *P* < 0.001). However, the difference in *P. carotovorum* growth rate between wild type and *PpRALF1* KO plants was not significant (Figure 4C). The growth of *F. solani* was significantly reduced in all *PpRALF* knockouts in comparison to wild type plants (Figure 4C; ANOVA with post-hoc Tukey HSD *P* < 0.001). Herewith, *PpRALF2* KO and *PpRALF3* KO mutant lines were the most resistant to *F. solani* infection. Taken together, these results indicated that knockout of *PpRALF* genes increased resistance of *P. patens* to phytopathogens. Apparently, the PpRALF2 and PpRALF3 play a key role in negative regulation of immune response in *P. patens* as was shown for some members of the RALF family in Arabidopsis.

### 3.5 Transcriptome response during *F. solani* infection in *PpRALF3* KO line

According to our analysis, *PpRALF2* and *3* KO genotypes were more resistant to phytopathogens than wild type plants and *PpRALF1* knockout lines. To expand our understanding of the roles of PpRALFs in immune stress response, we next sought to perform RNA sequencing (RNA-seq) analysis to compare transcriptome response of *PpRALF3* KO and wild type plants during *F. solani* infection. Twelve paired-end RNA-Seq libraries were generated from three biological replicates of mock-inoculated and *F. solani*-infected wild type and *PpRALF3* KO plants (Supplementary Table S3). Overall, we detected more than 1700 differentially regulated genes (DEGs; −1 ≤ log2 fold change ≥ 1, *Padj* < 0.05) in both genotypes and 474 DEGs were commonly regulated in wild type and knockout plants (Figure 5A; Supplementary Table S4). These DEGs were changed in a similar manner in both wild type and knockout plants (Figure 5B). We further used the g:Profiler (Kolberg et al., 2020) for finding enriched GO terms in the set of commonly regulated DEGs (Figure 5C). These DEGs were mainly enriched in such GO terms as oxidoreductase activity (GO:0016491), response to reactive oxygen species (GO:0000302), antioxidant activity (GO:0016209; Supplementary Table S5). These GO terms include genes involved in detoxification of oxygen radicals, such as superoxide dismutase (Pp3c17_14510) or superoxide dismutase 1 copper chaperone (Pp3c17_14500) and producing lignin-like compounds, such as cinnamyl alcohol dehydrogenases (Pp3c7_12220). These findings corroborate previous data obtained during transcriptome analysis of the interaction between *Physcomitrium patens* and *Botrytis cinerea* (Reboledo et al., 2021). We also identified some members of the EXPANSIN protein family, such as Pp3c11_12000 (expansin 5-related) and Pp3c7_12870 (expansin 5-related) that were upregulated in infected plants.

**Figure 5.**
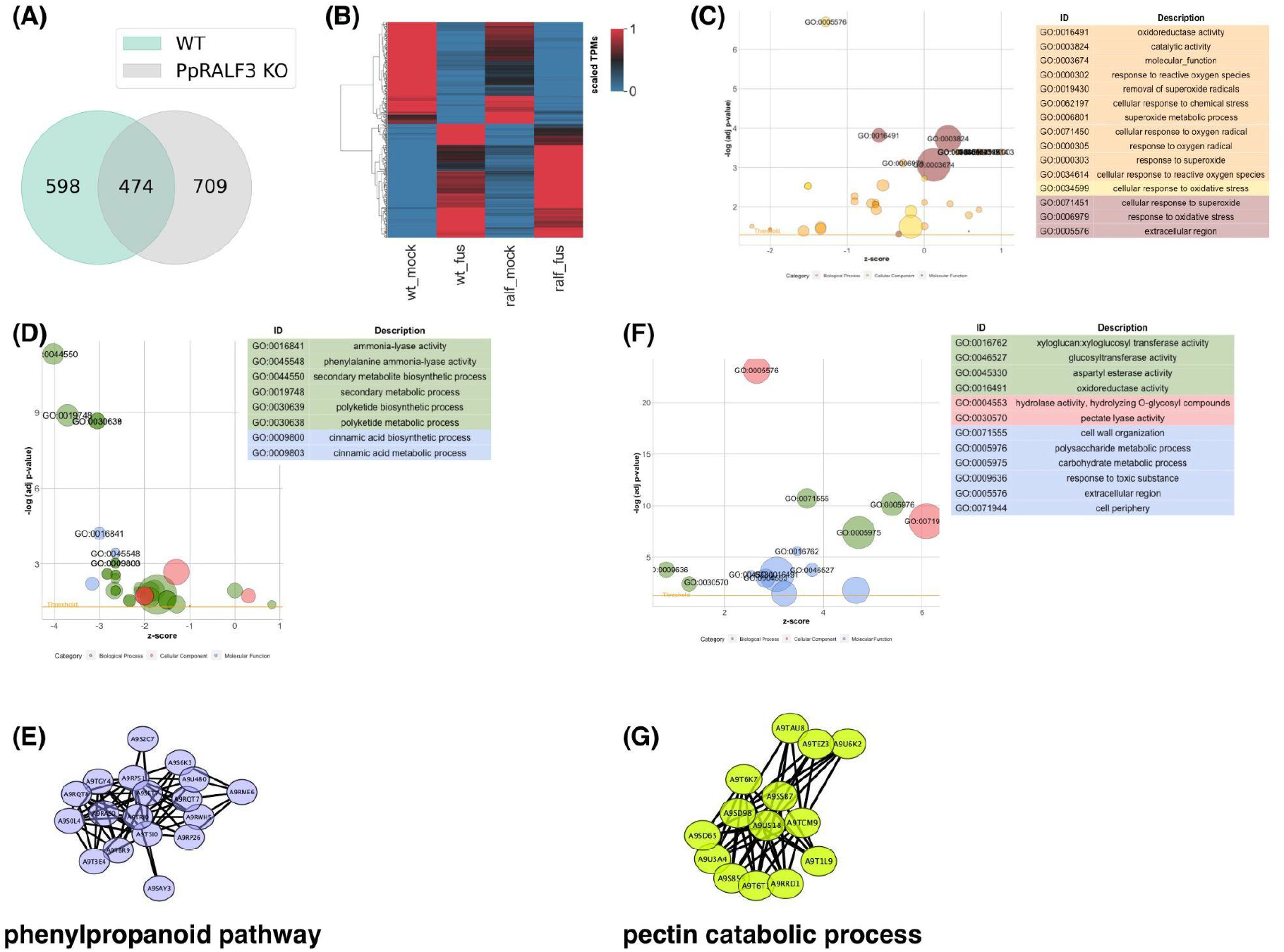
(**A)** - Venn diagram shows overlapping differentially regulated genes (DEGs; 1 ≤ log2 fold change ≥ 1, *Padj* < 0.05) between wild type and *PpRALF3* KO lines; **(B)** - Heatmap that depicts the euclidean-distance clustering analysis of overlapped DEGs. wt_mock - scaled mean TPM in wild type mock plants, wt_fus - scaled mean TPM in wild type infected plants, ralf_mock - scaled mean TPM in *PpRALF3* KO mock plants, ralf_fus - scaled mean TPM in *PpRALF3* KO infected plants; **(C)** - A bubble plot that depicts enriched GO terms (false discovery rate (FDR) adjusted *P*-values ≤ 0.05) for overlapped DEGs between wild type and *PpRALF3* KO lines; **(D)** - A bubble plot that depicts enriched GO terms in wild type plants. −Log adj p-values are shown in y-axis and z-scores are shown in x-axis. Size of the bubbles is proportional to the number of DEGs; **(E)** - Protein–protein interaction networks of a group of upregulated differentially expressed genes in wild type plants that belong to phenylpropanoid metabolic process. Genes are indicated with nodes, and interactions between proteins are represented by edges; **(F)** - A bubble plot that depicts enriched GO terms in *PpRALF3* KO plants. −Log adj p-values are shown in y-axis and z-scores are shown in x-axis. Size of the bubbles is proportional to the number of DEGs; **(G)** - Protein–protein interaction networks of a group of downregulated differentially expressed genes in *PpRALF3* KO plants that belong to pectin catabolic process. Genes are indicated with nodes, and interactions between proteins are represented by edges.

We next analyzed the differences in transcriptomic responses between examined genotypes. In wild type plants, most of the top ten significantly enriched GO terms belonged to secondary metabolite biosynthetic processes, including polyketide metabolic process (GO:0030638), phenylpropanoid metabolic process (GO:0009698), cinnamic acid metabolic process (GO:0009803; Figure 5D). The upregulated DEGs belonged to these GO terms included histidine and phenylalanine ammonia-lyases (e.g. Pp3c13_9000) and chalcone synthases (e.g. Pp3c11_9040, Figure 5F). In addition, the transcriptional level of genes encoding heat shock proteins increased in wild type plants. The downregulated DEGs identified in wild type plants were enriched in photosynthesis (GO:0015979). This is in line with the previous study on the interaction between *P. patens* and *B. cinerea* (Reboledo et al., 2021).

In contrast, the GO terms of downregulated genes in *PpRALF3* knockout line were predominantly enriched in processes related to cell wall organization and biogenesis, such as pectin catabolic process (GO:0045490), xyloglucan metabolic process (GO:0010411), polysaccharide metabolic process (GO:0005976; Figure 5F). For example, groups of pectate lyases (e.g. Pp3c21_16990) and pectin esterases were identified in this set of DEGs (Figure G). The upregulated DEGs were enriched by such GO terms as oxidoreductase activity (GO:0016491) and cell wall organization or biogenesis (GO:0071554). The latter included pectin esterases, different glycosyl hydrolases and expansins.

In conclusion, transcriptomic changes in wild type plants are in line with previous data on *P. patens* response to phytopathogens (Reboledo et al., 2021) and alter such processes as ROS production and detoxification, biosynthesis of secondary metabolites with different roles in defense and some others. The remarkable difference between *PpRALF3* KO and wild type plants consisted of transcriptomic changes in genes related to cell wall modification and biogenesis processes. Considering the known role of RALF peptides in cell wall remodeling, it can be suggested that these changes during infection result from knockout of *PpRALF3* gene.

## 4 Discussion

RAPID ALKALINIZATION FACTOR (RALF) peptides are ubiquitous for land plants, including *Physcomitrium patens* and *Selaginélla moellendorffii*, but were not identified in chlorophyte species (Campbell and Turner, 2017). In angiosperms, RALFs are shown to modulate responses to biotic and abiotic stresses (Blackburn et al., 2020), but it is currently unknown if ancient RALF peptides helped plants to cope with different stress factors or if the immune-related RALFs appeared later as a result of tandem duplication and diversification (Cao and Shi, 2012). Here, we investigated the role of three PpRALF peptides from the model bryophyte *P. patens*, named as PpRALF1 (Pp3c3_15280), PpRALF2 (Pp3c6_7200) and PpRALF3 (Pp3c25_4180) in the immune response. Previously, PpRALF1 and PpRALF2 have been shown to promote protonema tip growth and elongation (Ginanjar et al., 2022).

According to our findings, the knockout of all three *PpRALF* genes led to consistent changes in plant proteomes, suggesting that all three PpRALFs are functional. For example, a predicted metalloproteinase Pp3c15_830 was significantly downregulated in all *PpRALF* knockout lines. Plant metalloproteinases are shown to be involved in growth, development, and immunity (Flinn, 2008; Mishra et al., 2021).

The role of RALF peptides in stress response is mainly explored in the context of Arabidopsis biology. In Arabidopsis, several RALF peptides have shown the ability to modulate elf18-induced ROS production (Abarca et al., 2021). Based the amino acid compositions, we clustered PpRALFs with AtRALFs, such as AtRALF23 and AtRALF33, that negatively regulate immune response and inhibit elf18-induced ROS production in Arabidopsis (Stegmann et al., 2017; Abarca et al., 2021). In addition, the AtRALF23 and AtRALF33 were shown to interfere PAMP-triggered immunity (PTI) by binding to receptor FER-LLG1 complex and inhibition of its scaffold function (Shen et al., 2017; Rzemieniewski and Stegmann, 2022). AtRALF23 overexpression increases susceptibility to *Pseudomonas syringae* pv. *tomato* DC3000 and to the fungus *Plectosphaerella cucumerina* (Stegmann et al., 2017) and LRX3/4/5-RALF22/23-FER module negatively regulates the levels of jasmonic acid (JA), salicylic acid (SA) and abscisic acid (ABA) in Arabidopsis (Zhao et al., 2021). The hallmark of AtRALFs, that negatively regulate immune response, is the dibasic site “RR” for subtilase S1P (Srivastava et al., 2009; Stegmann et al., 2017; Abarca et al., 2021). However, there are some exceptions from this rule in Arabidopsis. In *P. patens*, only PpRALF1 protein precursor contains the corresponding “RRLL” motif (Ginanjar et al., 2022). According to our data, the PpRALF1 peptide has no role in stress response, but the knockouts of two *PpRALFs* (2 and 3) resulted in increasing resistance to bacterial and fungal pathogens - *P. carotovorum* and *F. solani*, suggesting the negative regulation of immune response by these peptides. These PpRALFs are also placed in a distinct group on the phylogenetic tree (Ginanjar et al., 2022).

In land plants, the immune response resulted in extensive transcriptome reprogramming (Bjornson et al., 2021; Campos et al., 2021; Reboledo et al., 2021). The early transcriptome changes include upregulation of genes involved in response to chitin and wounding at 5 min and participated in response to hydrogen peroxide and cell wall modification at 180 min after treatment by known elicitors in the model plant Arabidopsis (Bjornson et al., 2021). In our study, we found that transcriptome reprogramming in wild type *P. patens* plants at 7 dpi affected genes that participate in response to reactive oxygen species (GO:0000302), polyketide metabolic process (GO:0030638), phenylpropanoid metabolic process (GO:0009698), cinnamic acid metabolic process (GO:0009803). It has been shown that ROS level rises quickly in *P. patens* cells during *B. cinerea* infection and plays a role as second messengers during the moss immunity (Ponce de León et al. 2012). In addition, the moss *P. patens* responds to the fungal pathogen *B. cinerea* by reinforces the cell wall, upregulating the genes involved in defense response and activation of the shikimate and phenylpropanoid pathways (Ponce De León et al., 2012; Castro et al., 2016; Reboledo et al., 2021). The *Marchantia* response to oomycete infection is also based on the phenylpropanoid-mediated biochemical defenses that suggest this mechanism as a hallmark of an ancestral pathogen deterrence strategy (Carella et al., 2019).

Even though the difference in the pathogenic agent - *B. cinerea* (necrotrophic fungus) vs *F. solani* (facultative parasite) and time after inoculation when samples were collected - 24-h vs 7 dpi, about 40% of DEGs in wild type plants from our data were identical to the Reboledo dataset (Reboledo et al., 2021). However, the overlap between DEGs from *PpRALF3* KO plants and the aforementioned dataset was only 27%, suggesting specific immune responses in the knockout line.

Because our ultimate goal was to understand the role of PpRALFs in stress response, we concentrated on the comparison of the transcriptomes between wild type and *PpRALF3* knockout plants during *F. solani* infection. About 40% of *PpRALF3* KO DEGs were identical to wild type DEGs and changed in similar manner. The corresponding GO terms belonged to oxidoreductase activity (GO:0016491), response to reactive oxygen species (GO:0000302), antioxidant activity (GO:0016209). This finding is in line with previous studies on vascular and non-vascular plants in which increased expression of oxidative stress related genes encoding peroxiredoxins, thioredoxins, ferredoxins during infection was detected (Ponce de León, 2011; Bjornson et al., 2021; Reboledo et al., 2021). According to our results, the *PpRALF3* KO DEGs were not enriched in phenylpropanoid and cinnamic acid metabolic processes as we observed in wild type plants, but some genes involved in the flavonoid biosynthetic process, such as chalcone synthases, were upregulated in mutant lines. Importantly, the *PpRALF2* KO and *PpRALF3* KO lines were less infected by *F. solani* than wild type plants and *PpRALF1* KO genotype suggest their increased resistance. Therefore, the transcriptome differences between wild type and *PpRALF3* KO plants might reflect reduced propagation of *F. solani* in knockout lines. In contrast to wild type plants, a group of genes involved in the pectin catabolic process, such as pectate lyases and pectin esterases, were significantly downregulated in the *PpRALF3* KO plants under infection. Pectins are structural heteropolysaccharides and major components of the plant primary cell wall involved in maintaining plant growth and development, morphogenesis, defense responses, etc. (Hongo et al., 2012; Leng et al., 2017). The pectin-degrading enzymes cause plant tissue maceration, cell lysis and modification of the cell wall structure (Atanasova et al., 2018). Our findings is corroborate with the previous findings that salt stress resulted in downregulation of genes involved in cell wall organization and modification processes, such as pectin lyase-like superfamily proteins, expansins, xyloglucan hydrolases in *lrx345* (lrx3, 4, 5 triple mutants) Arabidopsis plants (Zhao et al., 2021). LRX8-LRX11 proteins are shown to interact with RALF4/19 and regulate pollen germination and pollen tube growth in Arabidopsis (Mecchia et al., 2017). The downregulation of similar cell wall related genes in our *PpRALF3* knockout line under stress conditions suggest the role of RALF peptides and the cognate receptors, such as FERONIA or LRXs, as modules that integrate plant growth and stress tolerance regulation in land plants.

In addition, it has been shown that pectin methylesterification is a subject of regulation during response to phytopathogens and a high level of pectin methylesterification correlated with an increased resistance to pathogens (Wydra and Beri, 2006; Lionetti et al., 2012). In Arabidopsis, downregulation of some pectin methylesterases during *B. cinerea* infection probably represent a defense mechanism to limit pectin demethylesterification and its subsequent degradation by fungal enzymes (Lionetti et al., 2017). Together with our results on the increased cell wall regeneration in *PpRALF2* and *PpRALF3* KO protoplasts relative to wild type cells, it may point to changes in cell wall composition as an important factor of increased resistance to phytopathogens in the corresponding knockout lines. There is no direct evidence how different RALF peptides influence cell wall composition in plants, but knockout of some LRXs in Arabidopsis resulted in the increase of mannose and lignin if compared with wild type plants (Draeger et al., 2015). Taken together, our findings suggest the role of PpRALF2 and PpRALF3 peptides in negative regulation of *P. patens* immune response.

We observed that *PpRALF3* and both double KO mutant lines were more tolerant to growth inhibition under salt and oxidative stress conditions. Previously, the role of AtRALF22 or AtRALF23 and their cognate receptors in coordinated regulation of cell wall integrity, growth and salt stress response was demonstrated in Arabidopsis (Zhao et al., 2018, 2021). The overexpression of AtRALF22 or AtRALF23 are shown to increase sensitivity to salt stress (Zhao et al., 2018). Moreover, LRX3/4/5-RALF22/23-FER module is shown to regulate hormonal homeostasis and ROS accumulation in Arabidopsis. Thus, our findings are in line with previous studies on flowering plants and show that RALF peptides in the non-vascular plants can also participate in abiotic stress response, modulating plant growth in such conditions. The detailed mechanisms through which ancient RALF and their cognate receptors coordinated plant growth, cell wall integrity and response to distinct environmental changes (e.g., pathogen invasion) is unknown. However, the future study on genetically redundant plants, such as *P. patens*, can help elucidate the evolution and exact mechanisms of RALF peptides signaling.

## 5 Data availability statement

The nucleotide sequence reported in this paper has been submitted to NCBI Sequence Reads Archive (SRA) with accession number PRJNA879762. The mass spectrometry proteomics data have been deposited to the ProteomeXchange Consortium via the PRIDE (Perez-Riverol et al., 2022) partner repository with the dataset identifier PXD037111.

**Password:** 7qCU3gkd

## 6 Author contributions

AM, IL, IF wrote the manuscript. AM, IL, NG conducted all experiments and contributed to data analysis. IF performed data analysis and supervised the study. AK, VL and DKh conducted generation of knockout moss lines, also AK contributed to growth and morphological experiments. TM, ECh, SE performed pathogens inoculation and moss treatment. VV, KK, TG and VB performed RNA-seq analysis.

## 9 Acknowledgments

We thank the Center for Precision Genome Editing and Genetic Technologies for Biomedicine, Federal Research and Clinical Center of Physical-Chemical Medicine of the Federal Medical Biological Agency for the expertise and guidance in genetic engineering.

## 10 Supplementary material

## References

Abarca, A., Franck, C. M., and Zipfel, C. (2021). Family-wide evaluation of RAPID ALKALINIZATION FACTOR peptides. Plant Physiol. 187, 996–1010.

Akita, M., Lehtonen, M. T., Koponen, H., Marttinen, E. M., and Valkonen, J. P. T. (2011). Infection of the Sunagoke moss panels with fungal pathogens hampers sustainable greening in urban environments. Sci. Total Environ. 409, 3166–3173.

Alvarez, A., Montesano, M., Schmelz, E., and Ponce de León, I. (2016). Activation of Shikimate, Phenylpropanoid, Oxylipins, and Auxin Pathways in Pectobacterium carotovorum Elicitors-Treated Moss. Front. Plant Sci. 7, 328.

Atanasova, L., Dubey, M., Grujic, M., Gudmundsson, M., Lorenz, C., Sandgren, M., et al. (2018). Evolution and functional characterization of pectate lyase PEL12, a member of a highly expanded Clonostachys rosea polysaccharide lyase 1 family. BMC Microbiol. 18, 178.

Atkinson, N. J., Lilley, C. J., and Urwin, P. E. (2013). Identification of genes involved in the response of Arabidopsis to simultaneous biotic and abiotic stresses. Plant Physiol. 162, 2028–2041.

Bjornson, M., Pimprikar, P., Nürnberger, T., and Zipfel, C. (2021). The transcriptional landscape of Arabidopsis thaliana pattern-triggered immunity. Nat Plants 7, 579–586.

Blackburn, M. R., Haruta, M., and Moura, D. S. (2020). Twenty Years of Progress in Physiological and Biochemical Investigation of RALF Peptides. Plant Physiol. 182, 1657–1666.

Bolger, A. M., Lohse, M., and Usadel, B. (2014). Trimmomatic: a flexible trimmer for Illumina sequence data. Bioinformatics 30, 2114–2120.

Campbell, L., and Turner, S. R. (2017). A Comprehensive Analysis of RALF Proteins in Green Plants Suggests There Are Two Distinct Functional Groups. Front. Plant Sci. 8, 37.

Campos, M. D., Félix, M. do R., Patanita, M., Materatski, P., and Varanda, C. (2021). High throughput sequencing unravels tomato-pathogen interactions towards a sustainable plant breeding. Hortic Res 8, 171.

Cao, J., and Shi, F. (2012). Evolution of the RALF Gene Family in Plants: Gene Duplication and Selection Patterns. Evol. Bioinform. Online 8, 271–292.

Carella, P., Gogleva, A., Hoey, D. J., Bridgen, A. J., Stolze, S. C., Nakagami, H., et al. (2019). Conserved Biochemical Defenses Underpin Host Responses to Oomycete Infection in an Early-Divergent Land Plant Lineage. Curr. Biol. 29, 2282–2294.e5.

Castro, A., Vidal, S., and Ponce de León, I. (2016). Moss Pathogenesis-Related-10 Protein Enhances Resistance to Pythium irregulare in Physcomitrella patens and Arabidopsis thaliana. Front. Plant Sci. 7, 580.

Chen, Z., Zhao, P., Li, F., Leier, A., Marquez-Lago, T. T., Wang, Y., et al. (2018). iFeature: a Python package and web server for features extraction and selection from protein and peptide sequences. Bioinformatics 34, 2499–2502.

Chevalier, E., Loubert-Hudon, A., and Matton, D. P. (2013). ScRALF3, a secreted RALF-like peptide involved in cell-cell communication between the sporophyte and the female gametophyte in a solanaceous species. Plant J. 73, 1019–1033.

Collonnier, C., Epert, A., Mara, K., Maclot, F., Guyon-Debast, A., Charlot, F., et al. (2017). CRISPR-Cas9-mediated efficient directed mutagenesis and RAD51-dependent and RAD51-independent gene targeting in the mossPhyscomitrella patens. Plant Biotechnol. J. 15, 122–131.

Draeger, C., Ndinyanka Fabrice, T., Gineau, E., Mouille, G., Kuhn, B. M., Moller, I., et al. (2015). Arabidopsis leucine-rich repeat extensin (LRX) proteins modify cell wall composition and influence plant growth. BMC Plant Biol. 15, 155.

Faurobert, M., Pelpoir, E., and Chaïb, J. (2007). Phenol extraction of proteins for proteomic studies of recalcitrant plant tissues. Methods Mol. Biol. 355, 9–14.

Fesenko, I. A., Arapidi, G. P., Skripnikov, A. Y., Alexeev, D. G., Kostryukova, E. S., Manolov, A. I., et al. (2015). Specific pools of endogenous peptides are present in gametophore, protonema, and protoplast cells of the moss Physcomitrella patens. BMC Plant Biol. 15, 87.

Fesenko, I., Kirov, I., Kniazev, A., Khazigaleeva, R., Lazarev, V., Kharlampieva, D., et al. (2019). Distinct types of short open reading frames are translated in plant cells. Genome Res. 29, 1464–1477.

Fesenko, I., Shabalina, S. A., Mamaeva, A., Knyazev, A., Glushkevich, A., Lyapina, I., et al. (2021a). A vast pool of lineage-specific microproteins encoded by long non-coding RNAs in plants. Nucleic Acids Res. 49, 10328–10346.

Fesenko, I., Spechenkova, N., Mamaeva, A., Makhotenko, A. V., Love, A. J., Kalinina, N. O., et al. (2021b). Role of the methionine cycle in the temperature-sensitive responses of potato plants to potato virus Y. Mol. Plant Pathol. 22, 77–91.

Flinn, B. S. (2008). Plant extracellular matrix metalloproteinases. Funct. Plant Biol. 35, 1183–1193.

Franck, C. M., Westermann, J., and Boisson-Dernier, A. (2018). Plant Malectin-Like Receptor Kinases: From Cell Wall Integrity to Immunity and Beyond. Annu. Rev. Plant Biol. 69, 301–328.

Frederick, R. O., Haruta, M., Tonelli, M., Lee, W., Cornilescu, G., Cornilescu, C. C., et al. (2019). Function and solution structure of the Arabidopsis thaliana RALF8 peptide. Protein Sci. 28, 1115–1126.

Ge, Z., Bergonci, T., Zhao, Y., Zou, Y., Du, S., Liu, M.-C., et al. (2017). Arabidopsis pollen tube integrity and sperm release are regulated by RALF-mediated signaling. Science 358, 1596–1600.

Ge, Z., Dresselhaus, T., and Qu, L.-J. (2019). How CrRLK1L Receptor Complexes Perceive RALF Signals. Trends Plant Sci. 24, 978–981.

Ginanjar, E. F., Teh, O.-K., and Fujita, T. (2022). Characterisation of rapid alkalinisation factors in Physcomitrium patens reveals functional conservation in tip growth. New Phytol. 233, 2442–2457.

Herrera-Ubaldo, H., and de Folter, S. (2018). Exploring Cell Wall Composition and Modifications During the Development of the Gynoecium Medial Domain in Arabidopsis. Front. Plant Sci. 9, 454.

Hoang, Q. T., Cho, S. H., McDaniel, S. F., Ok, S. H., Quatrano, R. S., and Shin, J. S. (2009). An actinoporin plays a key role in water stress in the moss Physcomitrella patens. New Phytol. 184, 502–510.

Hongo, S., Sato, K., Yokoyama, R., and Nishitani, K. (2012). Demethylesterification of the primary wall by PECTIN METHYLESTERASE35 provides mechanical support to the Arabidopsis stem. Plant Cell 24, 2624–2634.

Kabir, M. N., Taheri, A., and Dumenyo, C. K. (2020). Development of PCR-Based Detection System for Soft Rot Pectobacteriaceae Pathogens Using Molecular Signatures. Microorganisms 8. doi: 10.3390/microorganisms8030358.

Katoh, K., Rozewicki, J., and Yamada, K. D. (2019). MAFFT online service: multiple sequence alignment, interactive sequence choice and visualization. Brief. Bioinform. 20, 1160–1166.

Khraiwesh, B., Qudeimat, E., Thimma, M., Chaiboonchoe, A., Jijakli, K., Alzahmi, A., et al. (2015). Genome-wide expression analysis offers new insights into the origin and evolution of Physcomitrella patens stress response. Sci. Rep. 5, 17434.

Kim, D., Langmead, B., and Salzberg, S. L. (2015). HISAT: a fast spliced aligner with low memory requirements. Nat. Methods 12, 357–360.

Kolberg, L., Raudvere, U., Kuzmin, I., Vilo, J., and Peterson, H. (2020). gprofiler2 -- an R package for gene list functional enrichment analysis and namespace conversion toolset g:Profiler. F1000Res. 9. doi: 10.12688/f1000research.24956.2.

Lang, D., Ullrich, K. K., Murat, F., Fuchs, J., Jenkins, J., Haas, F. B., et al. (2018). The Physcomitrella patens chromosome-scale assembly reveals moss genome structure and evolution. Plant J. 93, 515–533.

Leng, Y., Yang, Y., Ren, D., Huang, L., Dai, L., Wang, Y., et al. (2017). A Rice PECTATE LYASE-LIKE Gene Is Required for Plant Growth and Leaf Senescence. Plant Physiol. 174, 1151–1166.

Liao, Y., Smyth, G. K., and Shi, W. (2014). featureCounts: an efficient general purpose program for assigning sequence reads to genomic features. Bioinformatics 30, 923–930.

Li, C., Yeh, F.-L., Cheung, A. Y., Duan, Q., Kita, D., Liu, M.-C., et al. (2015). Glycosylphosphatidylinositol-anchored proteins as chaperones and co-receptors for FERONIA receptor kinase signaling in Arabidopsis. Elife 4. doi: 10.7554/eLife.06587.

Li, H., Handsaker, B., Wysoker, A., Fennell, T., Ruan, J., Homer, N., et al. (2009). The Sequence Alignment/Map format and SAMtools. Bioinformatics 25, 2078–2079.

Lionetti, V., Cervone, F., and Bellincampi, D. (2012). Methyl esterification of pectin plays a role during plant–pathogen interactions and affects plant resistance to diseases. J. Plant Physiol. 169, 1623–1630.

Lionetti, V., Fabri, E., De Caroli, M., Hansen, A. R., Willats, W. G. T., Piro, G., et al. (2017). Three Pectin Methylesterase Inhibitors Protect Cell Wall Integrity for Arabidopsis Immunity to Botrytis. Plant Physiol. 173, 1844–1863.

Loubert-Hudon, A., Mazin, B. D., Chevalier, É., and Matton, D. P. (2020). The ScRALF3 secreted peptide is involved in sporophyte to gametophyte signalling and affects pollen mitosis I. Plant Biol. 22, 13–20.

Maksimov, N. M., Breigin, M. A., and Ermakov, I. P. (2016). Regulation of ion transport across the pollen tube plasmalemma by hydrogen peroxide. Cell tissue biol. 10, 69–75.

McKinney, W. (2012). Python for Data Analysis: Data Wrangling with Pandas, NumPy, and IPython. “O’Reilly Media, Inc.”

Mecchia, M. A., Santos-Fernandez, G., Duss, N. N., Somoza, S. C., Boisson-Dernier, A., Gagliardini, V., et al. (2017). RALF4/19 peptides interact with LRX proteins to control pollen tube growth in Arabidopsis. Science 358, 1600–1603.

Merino, M. C., Guidarelli, M., Negrini, F., De Biase, D., Pession, A., and Baraldi, E. (2019). Induced expression of the Fragaria × ananassa Rapid alkalinization factor-33-like gene decreases anthracnose ontogenic resistance of unripe strawberry fruit stages. Mol. Plant Pathol. 20, 1252–1263.

Mishra, L. S., Kim, S.-Y., Caddell, D. F., Coleman-Derr, D., and Funk, C. (2021). Loss of Arabidopsis matrix metalloproteinase-5 affects root development and root bacterial communities during drought stress. Physiol. Plant. 172, 1045–1058.

Moussu, S., Broyart, C., Santos-Fernandez, G., Augustin, S., Wehrle, S., Grossniklaus, U., et al. (2020). Structural basis for recognition of RALF peptides by LRX proteins during pollen tube growth. Proc. Natl. Acad. Sci. U. S. A. 117, 7494–7503.

Murphy, E., and De Smet, I. (2014). Understanding the RALF family: a tale of many species. Trends Plant Sci. 19, 664–671.

Olsson, V., Joos, L., Zhu, S., Gevaert, K., Butenko, M. A., and De Smet, I. (2019). Look Closely, the Beautiful May Be Small: Precursor-Derived Peptides in Plants. Annu. Rev. Plant Biol. 70, 153–186.

Ow, S. Y., Salim, M., Noirel, J., Evans, C., Rehman, I., and Wright, P. C. (2009). iTRAQ Underestimation in Simple and Complex Mixtures: “The Good, the Bad and the Ugly.” J. Proteome Res. 8, 5347–5355.

Perez-Riverol, Y., Bai, J., Bandla, C., García-Seisdedos, D., Hewapathirana, S., Kamatchinathan, S., et al. (2022). The PRIDE database resources in 2022: a hub for mass spectrometry-based proteomics evidences. Nucleic Acids Res. 50, D543–D552.

Ponce de León, I. (2011). The Moss Physcomitrella patens as a Model System to Study Interactions between Plants and Phytopathogenic Fungi and Oomycetes. J. Pathog. 2011, 719873.

Ponce De León, I., Schmelz, E. A., Gaggero, C., Castro, A., Álvarez, A., and Montesano, M. (2012). Physcomitrella patens activates reinforcement of the cell wall, programmed cell death and accumulation of evolutionary conserved defence signals, such as salicylic acid and 12-oxo-phytodienoic acid, but not jasmonic acid, upon Botrytis cinerea infection. Mol. Plant Pathol. 13, 960–974.

Raudvere, U., Kolberg, L., Kuzmin, I., Arak, T., Adler, P., Peterson, H., et al. (2019). g:Profiler: a web server for functional enrichment analysis and conversions of gene lists (2019 update). Nucleic Acids Res. 47, W191–W198.

Reboledo, G., Agorio, A. D., Vignale, L., Batista-García, R. A., and Ponce De León, I. (2021). Transcriptional profiling reveals conserved and species-specific plant defense responses during the interaction of Physcomitrium patens with Botrytis cinerea. Plant Mol. Biol. 107, 365–385.

Robinson, M. D., McCarthy, D. J., and Smyth, G. K. (2010). edgeR: a Bioconductor package for differential expression analysis of digital gene expression data. Bioinformatics 26, 139–140.

Rzemieniewski, J., and Stegmann, M. (2022). Regulation of pattern-triggered immunity and growth by phytocytokines. Curr. Opin. Plant Biol. 68, 102230.

Schaefer, D. G., and Zrÿd, J. P. (1997). Efficient gene targeting in the moss Physcomitrella patens. Plant J. 11, 1195–1206.

Schessner, J. P., Voytik, E., and Bludau, I. (2022). A practical guide to interpreting and generating bottom-up proteomics data visualizations. Proteomics 22, e2100103.

Shannon, P., Markiel, A., Ozier, O., Baliga, N. S., Wang, J. T., Ramage, D., et al. (2003). Cytoscape: a software environment for integrated models of biomolecular interaction networks. Genome Res. 13, 2498–2504.

Shen, Q., Bourdais, G., Pan, H., Robatzek, S., and Tang, D. (2017). Arabidopsis glycosylphosphatidylinositol-anchored protein LLG1 associates with and modulates FLS2 to regulate innate immunity. Proc. Natl. Acad. Sci. U. S. A. 114, 5749–5754.

Solis-Miranda, J., Fonseca-García, C., Nava, N., Pacheco, R., and Quinto, C. (2020). Genome-Wide Identification of the CrRLK1L Subfamily and Comparative Analysis of Its Role in the Legume-Rhizobia Symbiosis. Genes 11. doi: 10.3390/genes11070793.

Srivastava, R., Liu, J.-X., Guo, H., Yin, Y., and Howell, S. H. (2009). Regulation and processing of a plant peptide hormone, AtRALF23, in Arabidopsis. Plant J. 59, 930–939.

Stegmann, M., Monaghan, J., Smakowska-Luzan, E., Rovenich, H., Lehner, A., Holton, N., et al. (2017). The receptor kinase FER is a RALF-regulated scaffold controlling plant immune signaling. Science 355, 287–289.

Szklarczyk, D., Franceschini, A., Wyder, S., Forslund, K., Heller, D., Huerta-Cepas, J., et al. (2014). STRING v10: protein–protein interaction networks, integrated over the tree of life. Nucleic Acids Res. 43, D447–D452.

Tena, G. (2016). Immunity: RALF to the rescue. Nat Plants 2, 16067.

Thynne, E., Saur, I. M. L., Simbaqueba, J., Ogilvie, H. A., Gonzalez-Cendales, Y., Mead, O., et al. (2017). Fungal phytopathogens encode functional homologues of plant rapid alkalinization factor (RALF) peptides. Mol. Plant Pathol. 18, 811–824.

Van Rossum, G., and Drake, F. L., Jr (1995). Python tutorial: Centrum voor Wiskunde en Informatica Amsterdam. The Netherlands.

Vidali, L., and Bezanilla, M. (2012). Physcomitrella patens: a model for tip cell growth and differentiation. Curr. Opin. Plant Biol. 15, 625–631.

Virtanen, P., Gommers, R., Oliphant, T. E., Haberland, M., Reddy, T., Cournapeau, D., et al. (2020). SciPy 1.0: fundamental algorithms for scientific computing in Python. Nat. Methods 17, 261–272.

Waskom, M. (2021). seaborn: statistical data visualization. J. Open Source Softw. 6, 3021.

Waterhouse, A. M., Procter, J. B., Martin, D. M. A., Clamp, M., and Barton, G. J. (2009). Jalview Version 2--a multiple sequence alignment editor and analysis workbench. Bioinformatics 25, 1189–1191.

Wieghaus, A., Prüfer, D., and Schulze Gronover, C. (2019). Loss of function mutation of the Rapid Alkalinization Factor (RALF1)-like peptide in the dandelion Taraxacum koksaghyz entails a high-biomass taproot phenotype. PLoS One 14, e0217454.

Wood, A. K. M., Walker, C., Lee, W.-S., Urban, M., and Hammond-Kosack, K. E. (2020). Functional evaluation of a homologue of plant rapid alkalinisation factor (RALF) peptides in Fusarium graminearum. Fungal Biol. 124, 753–765.

Wydra, K., and Beri, H. (2006). Structural changes of homogalacturonan, rhamnogalacturonan I and arabinogalactan protein in xylem cell walls of tomato genotypes in reaction to Ralstonia solanacearum. Physiol. Mol. Plant Pathol. 68, 41–50.

Xiao, Y., Stegmann, M., Han, Z., DeFalco, T. A., Parys, K., Xu, L., et al. (2019). Mechanisms of RALF peptide perception by a heterotypic receptor complex. Nature 572, 270–274.

Zhang, X., Peng, H., Zhu, S., Xing, J., Li, X., Zhu, Z., et al. (2020a). Nematode-Encoded RALF Peptide Mimics Facilitate Parasitism of Plants through the FERONIA Receptor Kinase. Mol. Plant 13, 1434–1454.

Zhang, X., Yang, Z., Wu, D., and Yu, F. (2020b). RALF–FERONIA Signaling: Linking Plant Immune Response with Cell Growth. Plant Communications 1, 100084.

Zhao, C., Jiang, W., Zayed, O., Liu, X., Tang, K., Nie, W., et al. (2021). The LRXs-RALFs-FER module controls plant growth and salt stress responses by modulating multiple plant hormones. Natl Sci Rev 8, nwaa149.

Zhao, C., Zayed, O., Yu, Z., Jiang, W., Zhu, P., Hsu, C.-C., et al. (2018). Leucine-rich repeat extensin proteins regulate plant salt tolerance in Arabidopsis. Proc. Natl. Acad. Sci. U. S. A. 115, 13123–13128.

