## Supplementary Figures for "RALF peptides modulate immune response in the moss *Physcomitrium patens*"

**A**

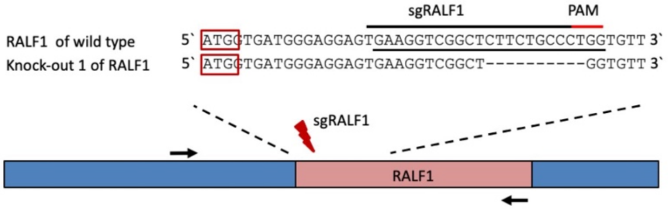


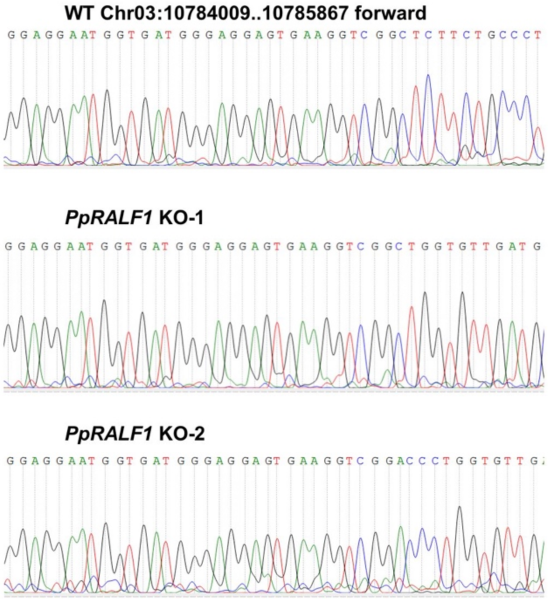


**Wild type of RALF1** - MVMGGVKVGSSALVLMLAMLAFVLVQCVDRADAGHG SPIMELATTSSGLPEGFDDLRMYEDPEDQFLDEPARRLLAARRRSYISYGALNRN RSPCPARSGRSYYTPNCNSNAGPARPYTRGCLRITRCQRV*

**Knock-out 1 of RALF1** - **MVMGGVKVG**WC*

**Knock-out 2 of RALF1** - **MVMGGVKVG**PWC*

**B**

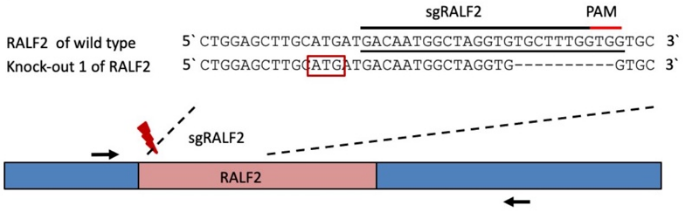


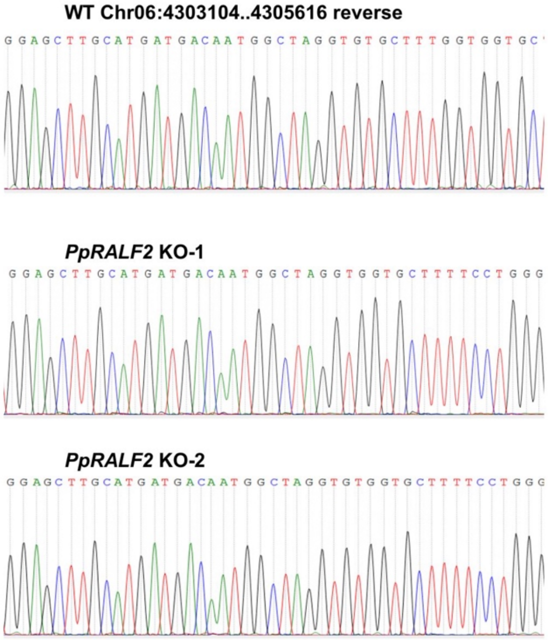


**Wild type of RALF2** -
MMTMARCALVVLFLGCLVLEAQAQAPFAAVAPSPFTGFPPQEAPAFAGSGGPVL YITYGALRANRSPCPAGAGRSYYTPNCGAASGPPNPYSRGCSYITRCARV*

**Knock-out 1 of RALF2** - MMTMARWCFSWVVWSWKPRRRLRSLP*

**Knock-out 2 of RALF2** – MMTMARCGAFPGLFGPGSPGAGSVRCRSSESLHRV PTAGGSRFRGLWWSGSLHHLRCSKSKPQSMPCWSRTKLLHAQLRCCIGTPEPI QQGLQLHHPLCSRLRRCTWVLPRSEMHMPSSTTMVE*

**C**

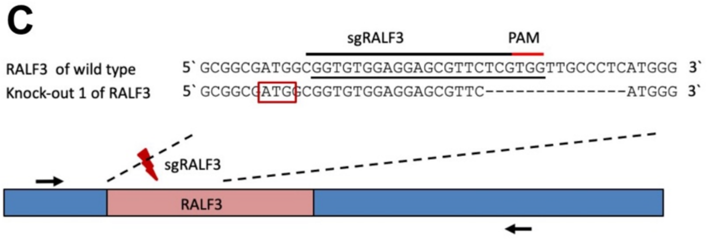


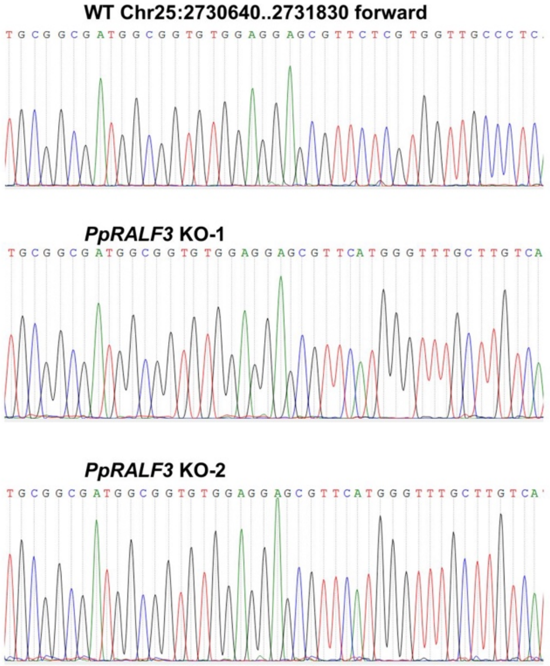


**Wild type of RALF3** -
MAVWRSVLVVALMGLLVMGIQAEQQLGFSPGAIYPDAHRVLAAAPNYYISYGSL
NADRAPCPASSSGRSYYTTNCQSSGTPQPYVRSCSQITRCARG*
**Knock-out 1 of RALF3** - MAVWRSVHGFACHGYPS*
**Knock-out 2 of RALF3** - MAVWRSVHGFACHGYPS*

**D**

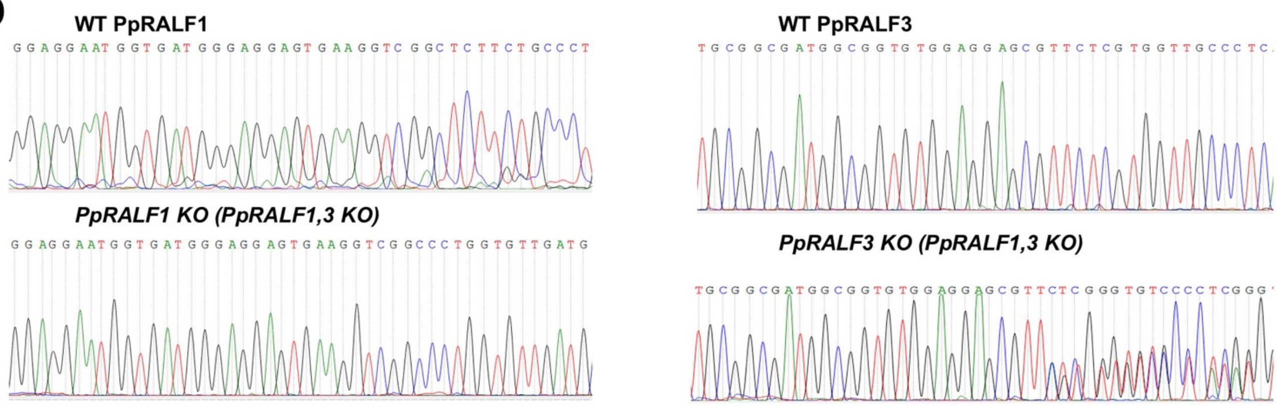


**Knock-out of RALF1 in PpRALF 1,3 KO** - MVMGGVKVGPGVDAGNAGIRAGAMRGQGGRGSWKPHHGTRYHVLRSPRGVR*
**Knock-out of RALF3 in PpRALF 1,3 KO** - MAVWRSVLGCPLGFARVGDTHSEPPCVFPPCAFYPYAPRVCGAAPI*

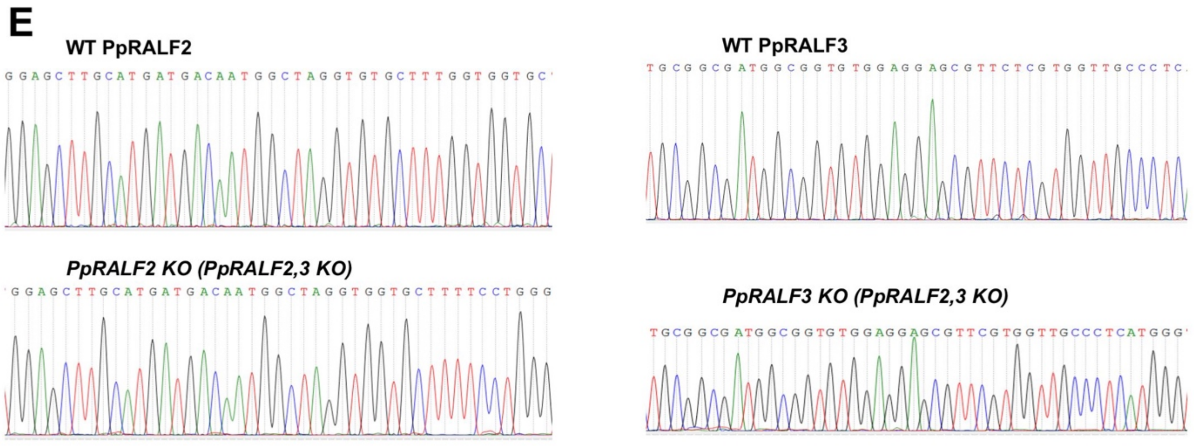


**Knock-out of RALF2 in PpRALF 2,3 KO** - MMTMARWCFSWVVWSWKPRRRLRSLP*
**Knock-out of RALF3 in PpRALF 2,3 KO** - MAVWRSVRGCPHGFACHGYPS*

**Supplementary Figure S1. A –** The schematic representation of sgRNAs designed to knockout of PpRALF1 peptide, the results of Sanger sequencing of obtained mutant lines and deduced amino acid sequences; **B –** The schematic representation of sgRNAs designed to knockout of PpRALF2 peptide, the results of Sanger sequencing of obtained mutant lines and deduced amino acid sequences; **C –** The schematic representation of sgRNAs designed to knockout of PpRALF3 peptide, the results of Sanger sequencing of obtained mutant lines and deduced amino acid sequences. *Black arrows* indicate the positions of primers for PCR analysis. *Red* frames indicate the positions of start codon; **D** - The results of sequencing of RALF genes in obtained mutant lines and deduced amino acid sequences for PpRALF1,3 double knockout; **E** - The results of sequencing of RALF genes in obtained mutant lines and deduced amino acid sequences for PpRALF2,3 double knockout

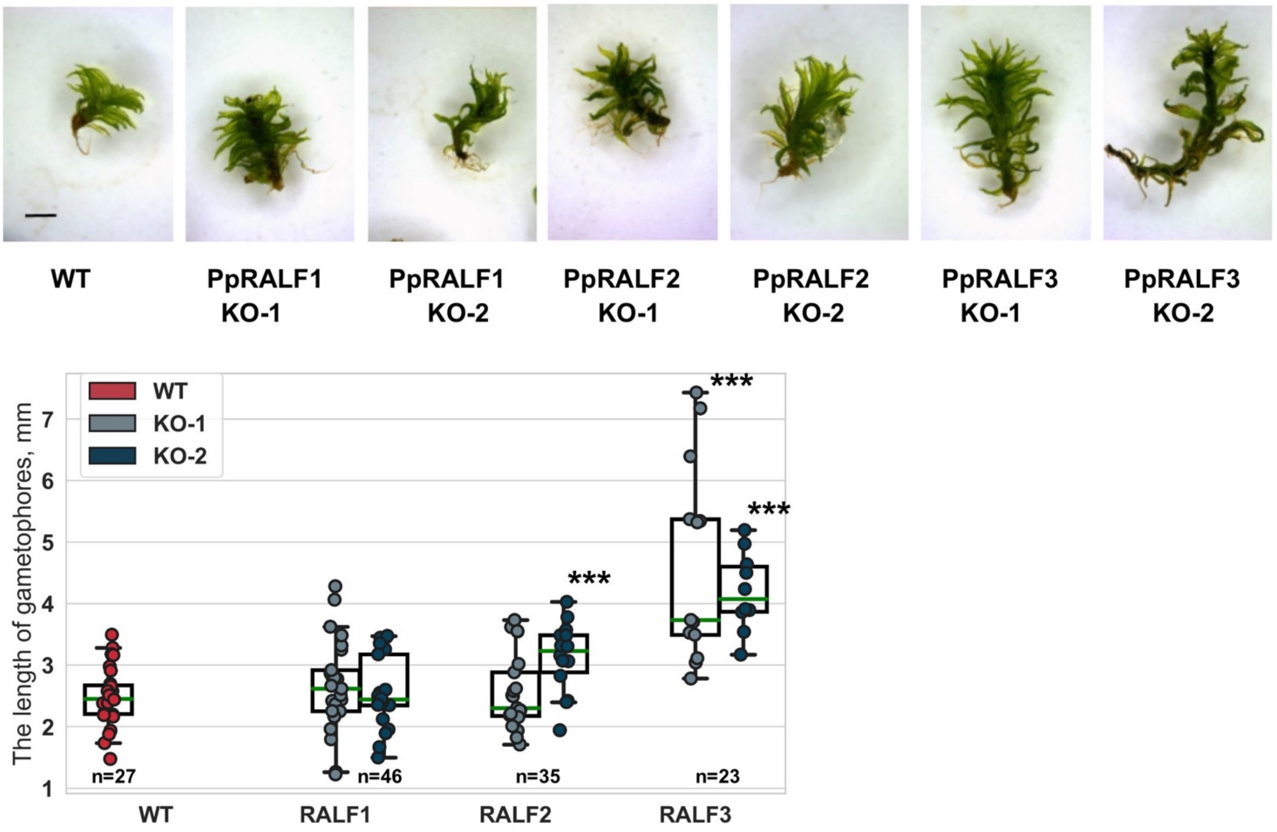


**Supplementary Figure S2.** The length of gametophores of wild-type and knockout plants. Analysis of variance and Tukey's HSD post hoc tests were performed ***$P < 0.001$

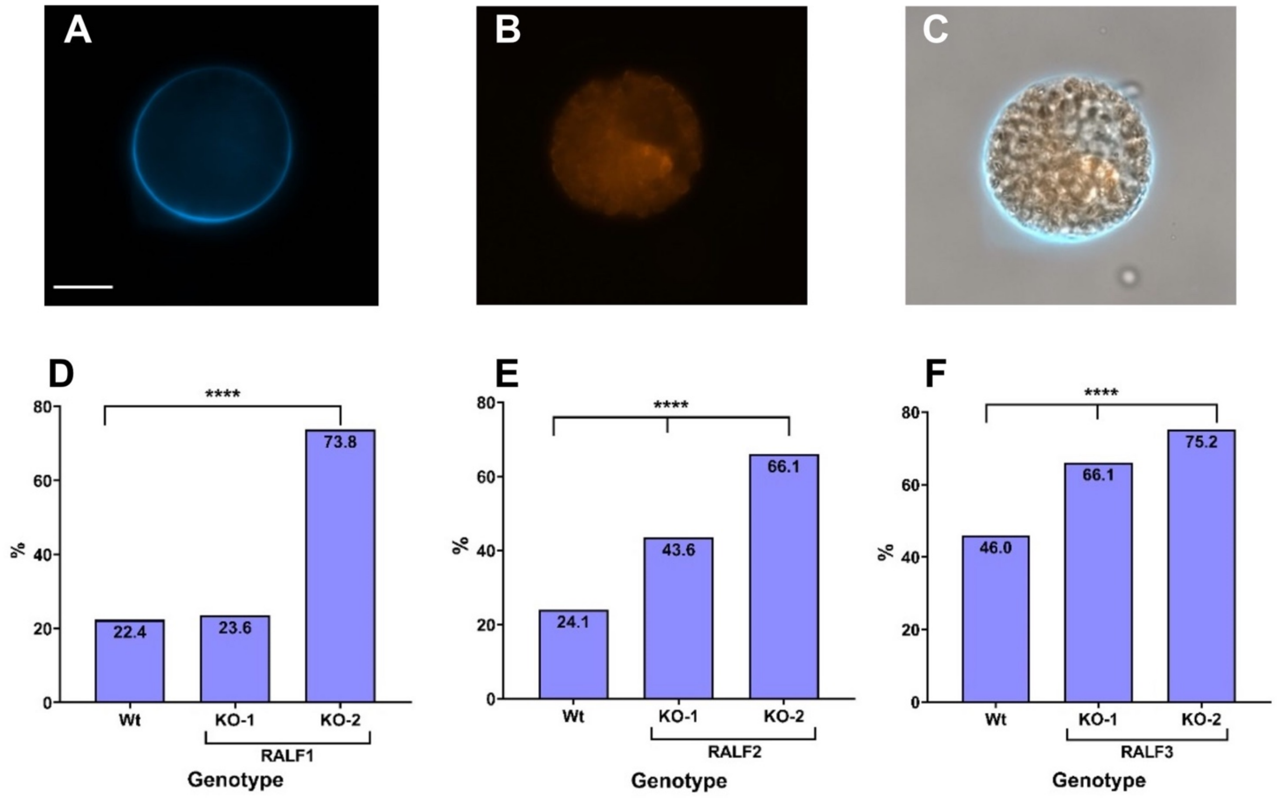


**Supplementary Figure S3.** **A** – Protoplast cell stained with Calcofluor White; **B** – Chloroplast autofluorescence in protoplast cell; **C** – combined A and B picture; **D** – The percent of regenerated protoplast in protoplast from PpRALF1 genotype in comparison to wild-type plants; **E** – The percent of regenerated protoplast in protoplast from PpRALF2 genotype in comparison to wild-type plants; **F** – The percent of regenerated protoplast in protoplast from PpRALF3 genotype in comparison to wild-type plants. Analysis of variance and Tukey's HSD post hoc tests were performed ****$P < 0.001$

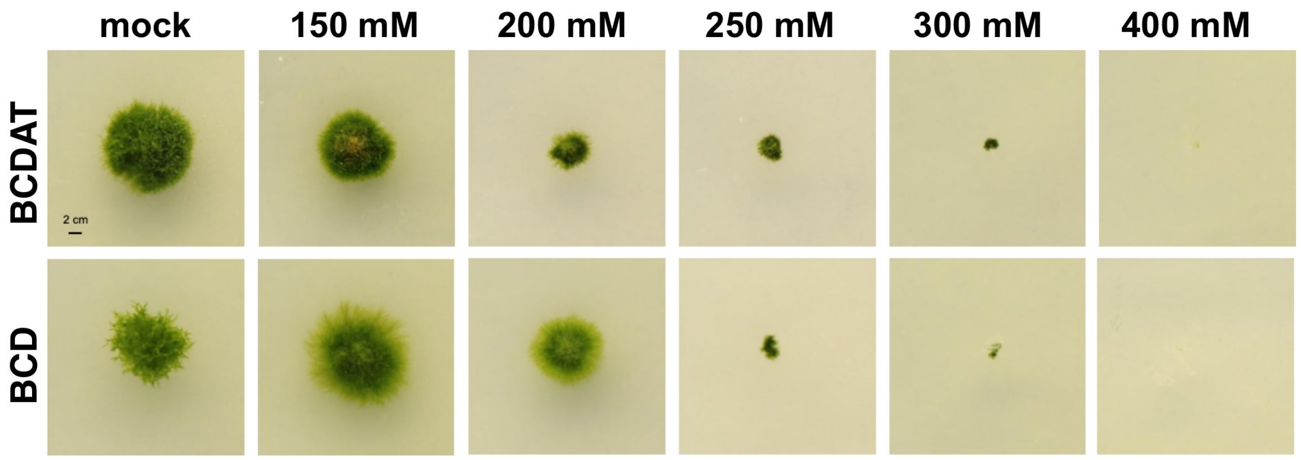


**Supplementary Figure S4.** The effect of different NaCl concentrations on growth rate in wild-type plants on two types of growing media: BCD and BCDAT (BCD medium supplemented with ammonium tartrate)
