## Supplementary Table S1 for "RALF peptides modulate immune response in the moss *Physcomitrium patens*"

**Supplementary Table S1.** List of primers used for creating genetic constructions and RT-PCR

| **Primer** | **Sequence** |
| --- | --- |
| ***Generating 20-mer guide sequence*** | |
| с3-1 top (RALF1) | CCATGAAGGTCGGCTCTTCTGCCC |
| c3-1 bottom (RALF1) | AAACGGGCAGAAGAGCCGACCTTC |
| c6-1 top (RALF2) | CCATGACAATGGCTAGGTGTGCTT |
| c6-1 bottom (RALF2) | AAACAAGCACACCTAGCCATTGTC |
| c25-1 top (RALF3) | CCATGGTGTGGAGGAGCGTTCTCG |
| c25-2 bottom (RALF3) | AAACCGAGAACGCTCCTCCACACC |
| ***Identification of clones with knockout RALF genes*** | |
| ra1f (RALF1) | TGGACGTAAGTGGTTTGGGT |
| ra1r (RALF1) | CAAGGGAGGACTCCGAAAGC |
| ra2f2 (RALF2) | GTTGTGTGGCTTCACTCGGA |
| ra2r2 (RALF2) | CCCTACCCTAGCCCAGACAA |
| ra3f1 (RALF3) | GGACTATTCCTGCTGCTGCG |
| ra3r1 (RALF1) | CGGCGTTCAGTGACCCATAG |
| ***RT-qPCR*** | |
| ACT_f | ACCGAGTCCAACATTCTACC |
| ACT_r | GTCCACATTAGATTCTCGCA |
| RALF1(Pp3c3_15280)_f | TGTTGATGCTGGCAATGCTG |
| RALF1(Pp3c3_15280)_r | CTACACTCTCTGGCAACGGG |
| RALF2(Pp3c6_7200)_f | TAAGAGCAAACCGCAGTCCA |
| RALF2(Pp3c6_7200)_r | GCAATTGGGCGTGTAGTAGC |
| RALF3(Pp3c25_4180)_f | GTTCTCGTGGTTGCCCTCAT |
| RALF3(Pp3c25_4180)_r | CAACGAGTGATCTGGGAGCA |
| PpEF_f | TTTGGGATTGAAATGTCGTG |
| PpEF_r | TGAGCATGAGAAATTGGGTCT |
| Pec3_f | GGGATTCGAAAAATTACTGGCTG |
| Pec3_r | GCTTTTCTTTCATCAACCCA |
| FS_f | TCGCGCCTTGATATTCCACA |
| FS_r | ACTTCCAGAGGGCAATGTCG |
